# A Dual-Site Inhibitor of CBP/p300 KIX is a Selective and Effective Modulator of Myb

**DOI:** 10.1101/2021.04.28.441843

**Authors:** Stephen T. Joy, Matthew J. Henley, Samantha N. De Salle, Matthew S. Beyersdorf, Isaac W. Vock, Allison J.L. Huldin, Anna K. Mapp

## Abstract

The protein-protein interaction between the KIX motif of the transcriptional coactivator CBP/p300 and the transcriptional activator Myb is a high value target due to its established role in certain acute myeloid leukemias (AML) and potential contributions to other cancers. However, the CBP/p300 KIX domain has multiple binding sites, several structural homologues, many binding partners, and substantial conformational plasticity, making it challenging to specifically target using small molecule inhibitors. Here, we report a picomolar dual-site inhibitor (MybLL-tide) of the Myb-CBP/p300 KIX interaction. MybLL-tide has higher affinity for CBP/p300 KIX than any previously reported compounds while also possessing 16,000-fold selectivity for the CBP/p300 KIX domain over other coactivator domains. MybLL-tide blocks the association of CBP and p300 with Myb in the context of the proteome leading to inhibition of key Myb•KIX-dependent genes in AML cells. These results show that MybLL-tide is an effective, modifiable tool to selectively target the KIX domain and assess transcriptional effects in AML cells and potentially other cancers featuring aberrant Myb behavior. Additionally, the dual-site design has applicability to the other challenging coactivators that bear multiple binding surfaces

---

CREB-binding protein (CBP) and p300 are large, homologous, multidomain coactivator proteins that perform a number of vital roles during transcription.^1,2^ These roles include regulating transcription and chromatin modification through a histone acetyltransferase domain, recruiting other transcriptional co-regulators, and localizing the site of transcription by interacting with DNA-bound transcriptional activators.^1–4^ CBP/p300 possesses at least four distinct activator-binding domains (ABDs) that interact with numerous different activators and other transcription factors, enabling CBP/p300 to control transcription of many genes but also requiring its ABDs to possess significant structural plasticity to accommodate the diversity of binding partners (Figure 1A).^5,6^ Notably, the CBP/p300 KIX domain is an ABD that interacts with a multitude of transcriptional activators, including Myb and MLL (Figure 1B),^6–11^ and a number of recent studies support an essential role of the Myb•KIX interaction in a subset of acute myeloid leukemias (AML).^12–16^ Abrogation of this interaction significantly attenuates AML growth and progression highlighting the Myb•KIX interaction as an important target for potential therapeutic intervention and study.^17–19^ Further studies have also shown Myb to play a vital role in metastasis and tumor progression in breast cancer, adenoid cystic carcinoma, and various other cancers, suggesting broad utility for potent inhibitors of Myb activity.^19–24^

**Figure 1:**
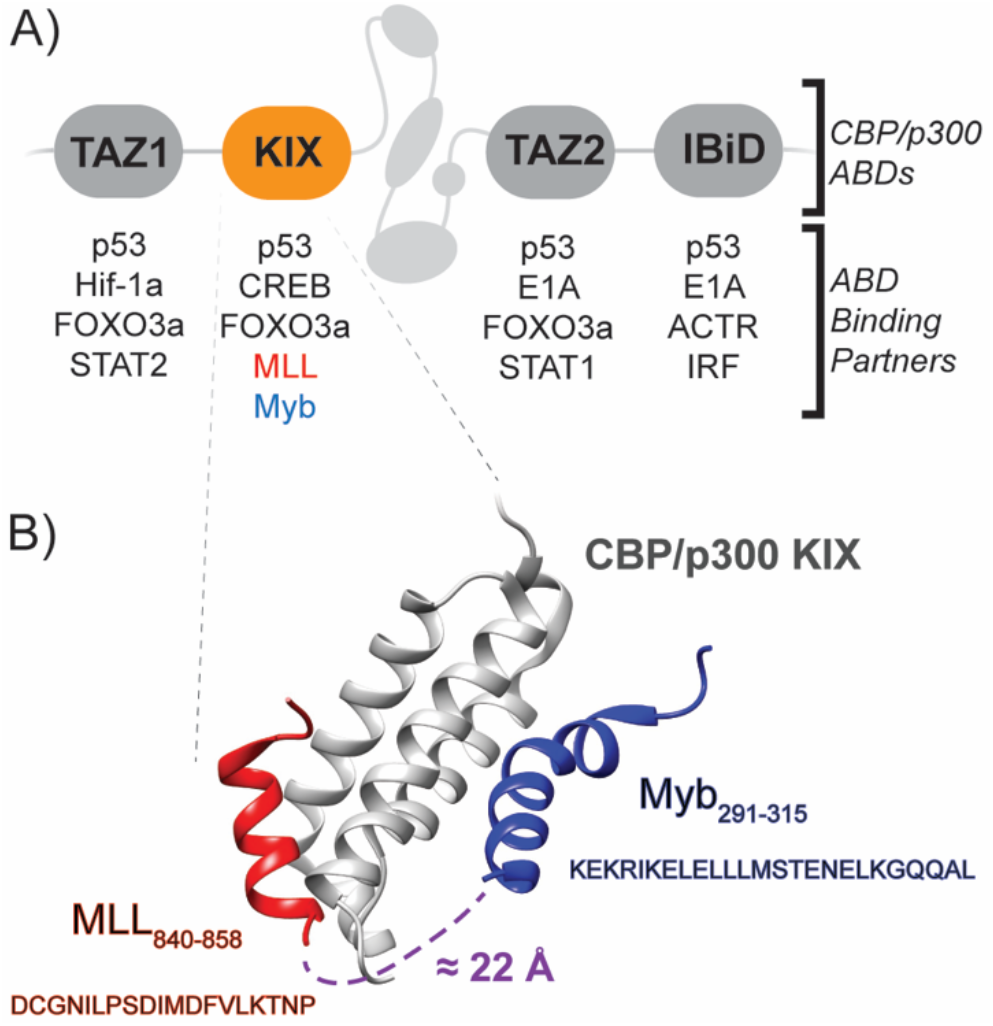
A) The four activator-binding domains of CBP that interact with various activators and other transcription factors through multiple binding sites per domain. Several transcription factors (p53, FOXO3a, and E1A) bind multiple CBP ABDs. B) The two distinct binding faces of the KIX domain, the MLL and Myb sites, the peptide sequences of the MLL (red) and Myb (blue) TADs, and the distance between the N-terminus of Myb and the C-terminus of MLL (purple). Schematic derived from protein data bank entry 2AGH.

The structural plasticity of the CBP/p300 KIX domain enables accommodation of many proteinogenic binding partners at two distinct binding sites (MLL and Myb sites, Figure 1B).^25^ Many of these partners, including the Myb transcriptional activation domain (TAD), have only modest affinity for KIX.^8,26,27^ KIX is also one of four ABDs within CBP/p300, all of which share binding partners (Figure 1A), and the KIX structural motif is present in a number of different proteins serving distinct roles.^28–32^ Altogether these characteristics make it particularly challenging to develop inhibitors that potently and selectively target CBP/p300 KIX protein-protein interactions.^33–38^

We hypothesized that linking the CBP/p300 KIX-binding TADs of Myb and MLL could produce a peptide with high affinity and selectivity for CBP/p300 KIX, even over structurally related coactivators. Targeting both binding sites simultaneously is likely to produce a higher affinity ligand than targeting the Myb face alone,^27^ and there are no other coactivators known that bind the Myb and MLL TADs simultaneously, suggesting a benefit for selectivity.^6,39–41^ Here we demonstrate that the resulting dual-site inhibitors, MybLL-tides, exhibit high affinity for CBP/p300 KIX (pM), >16,000-fold selectivity, and effectively block CBP and p300 from complexing with Myb. In doing so, MybLL-tide inhibits KIX•Myb-regulated genes in AML.

Analysis of existing NMR structural data for the ternary MLL•KIX•Myb complex indicated a distance of approximately 22 Å separating the C-terminus of MLL(840-858) and the N-terminus of Myb(291-315) (Figure 1B, Table S1).^25^ Based upon this, we designed three constructs in which the two peptides were connected via 1, 2, or 3 8-amino-3,6-dioxaoctanoic acid (AEEAc) monomers, chosen for their hydrophilicity, flexibility, and size (Table 1, compounds **1**-**3**). These peptide hybrids are designated as MybLL-tides and were successfully synthesized using automated peptide synthesis (See Supporting Information for details).

**Table 1.** MybLL-tides used in this study, N-terminal variations (X) and linker lengths (n). F852 (MLL) and L302 (Myb) residues are mutated to alanine for negative controls. Figure: Schematic of MybLL-tide constructs consisting of MLL(840-858) and Myb(291-315) bound by a flexible linker. Brackets flank AEEAc residue(s), dictating linker length. FITC = fluorescein isothiocyanate, Ac = Acetyl, TAT = Tat-cell penetrating peptide. GG = diglycine linker.

| MybLL | X | n | F852 | L302 |
| --- | --- | --- | --- | --- |
| **1** | **a:** FITC-β-Ala<br>**b:** Ac | 1 | F | L |
| **2** | **a:** FITC-β-Ala<br>**b:** Ac | 2 | F | L |
| **3** | **a:** FITC-β-Ala<br>**b:** Ac | 3 | F | L |
| **4** | **a:** FITC-β-Ala<br>**b:** Ac | 3 | A | A |
| **5** | Biotin-AEEAc | 3 | F | L |
| **6** | Biotin-AEEAc | 3 | A | A |
| **7** | FITC-β-Ala-TAT-GG | 3 | F | L |
| **8** | Ac-TAT-GG | 3 | F | L |
| **9** | Ac-TAT-GG | 3 | A | A |

Results from direct binding experiments with fluorescein-labeled versions of the three MybLL-tides (**1a, 2a, 3a**) indicate all three bind to CBP KIX with sufficiently high affinity such that the dissociation constants were beyond the measurement limits of the assay (<10 nM; Figure 2A, Table S2). Similar results were observed for p300 KIX (Figure S1), which has 90% sequence homology with CBP KIX and comparable affinity for Myb and MLL.^6^ Thermal stability experiments comparing unlabeled MybLL-tides **1b, 2b**, and **3b** in complex with CBP KIX showed a substantial increase in thermal stability relative to free CBP KIX for all three peptides with **1b** producing the least stable complex, but this assay also failed to determine whether **2b** or **3b** produced a more stable complex (Figure S2, Table S3). To differentiate between **2a** and **3a**, stopped-flow fluorescence binding experiments were employed to measure on and off-rates (Figure S3, Table S4), thus enabling calculation of dissociation constants and ultimately showing that the longest MybLL-tide, **3a**, has the highest affinity for CBP KIX, with a calculated dissociation constant of 360 ± 50 pM for CBP KIX (Figure 2B). In comparison, **2a** only possessed a K_D_ = 1.3 ± 0.08 nM for KIX, while the shortest MybLL-tide **1a** had the weakest K_D_ (11.3 ± 0.6 nM) (Figure 2B). Based on this cumulative data, all subsequent MybLL-tides incorporated three AEEAc units within the linker region. A negative control mutant (**4**, Table 1) was developed by introducing an F852A mutation on the MLL face and an L302A mutation on the Myb face. Both mutations are known to disrupt binding of the MLL and Myb TADs respectively,^12,34,43–46^ with **4a** exhibiting a >45,000-fold reduced affinity for KIX (Figure 2A). Subsequent Protein-Observed Fluorine (PrOF) NMR experiments further revealed that titrating unlabeled **3b** into a 3-fluorotyrosine-containing CBP KIX mutant produced ^19^F chemical shift perturbations at tyrosine residues in both the MLL and Myb sites,^42^ consistent with a model in which the high affinity of MybLL-ide **3** arises from interactions with both the Myb and MLL binding surfaces (Figure 2C, S4).

**Figure 2:**
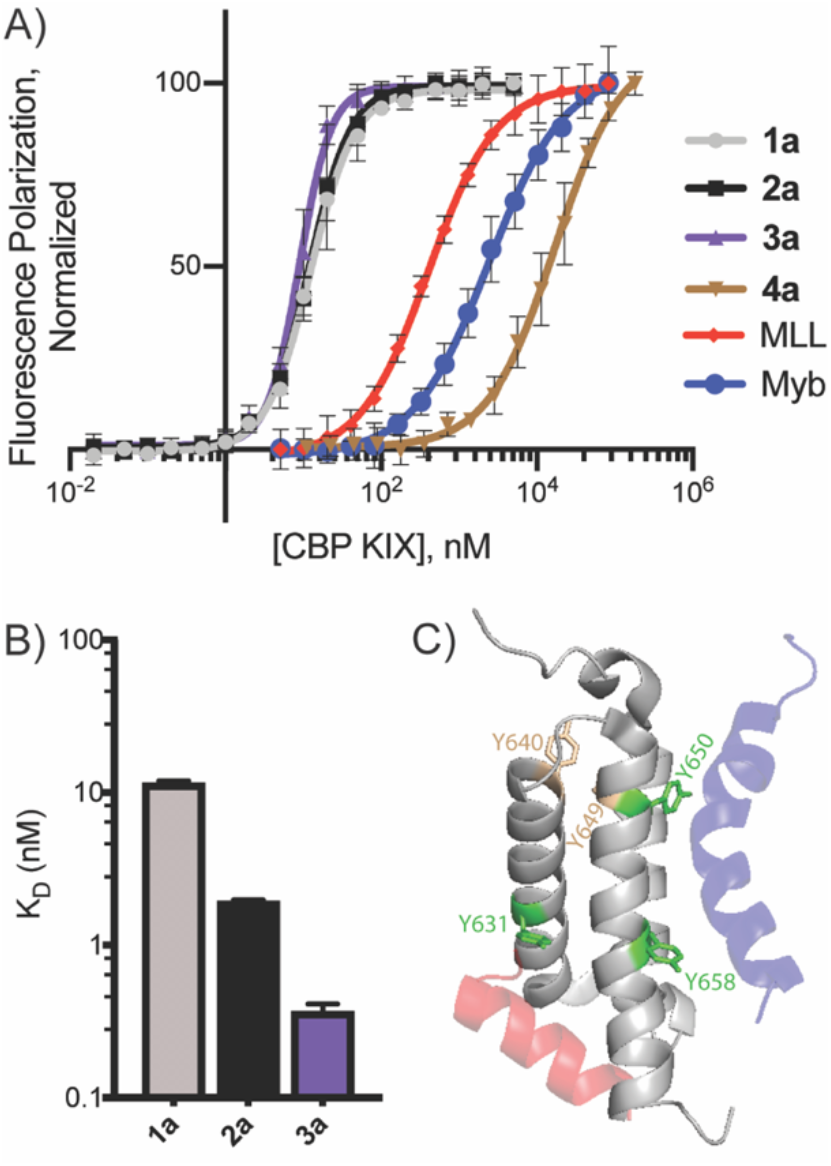
A) Direct-binding fluorescence polarization data of FITC-MybLL-tides, FITC-MLL, and FITC-Myb bound to CBP KIX. Error is SD derived from technical triplicates of experimental triplicates. B) Dissociation constants calculated from stopped-flow experiments on FITC-MybLL-tides with varying linker lengths. Error is SD of triplicate association experiments and 10-15 dissociation experiments. C) PrOF NMR experiment using 3-fluorotyrosine in place of native tyrosine residues on CBP KIX with shifting 3-fluorotyrosines labeled in green and non-responsive residues in tan. Titration of **3a** produces shifts of the two tyrosine residues on the Myb (blue) face of KIX (Y650, Y658) and the single tyrosine on the MLL (red) face of KIX (Y631). The two tyrosine residues that do not occupy either binding face do not shift significantly (Y640, Y649).

Because the CBP and p300 KIX motifs are the only known coactivators that form ternary complexes with Myb and MLL TADs, MybLL-tide’s enhanced affinity for CBP/p300 KIX was expected to yield improved selectivity CBP/p300 KIX over other related domains. To test this, a panel of coactivators including other CBP domains, KIX domains from other coactivators, and other MLL or Myb-binding proteins were assessed for **3a** affinity using fluorescence polarization (Figures 3A, S5). MybLL-tide **3a** was shown to have low affinity for several other CBP ABDs (TAZ1, IBiD) that also have flexible architecture, overlapping binding partners, and multiple binding sites.^47,48^ Additionally, **3a** exhibited low or unmeasurable affinity for other KIX domains, an important distinction as KIX is a common structural motif appearing in a variety of different proteins across different organisms (Figure 3A).^28–30^ In contrast, the MLL peptide interacts with many of these same coactivators with comparable affinity (µM dissociation constants, Table S5), while having 1,100-fold weaker affinity for CBP KIX relative to **3a** (Figure 3B).^49^ Ultimately, MybLL-tide’s enhanced affinity for CBP/p300 KIX results in 16,000-fold selectivity for the CBP/p300 KIX domain over other coactivator domains (Figure 3B).

**Figure 3:**
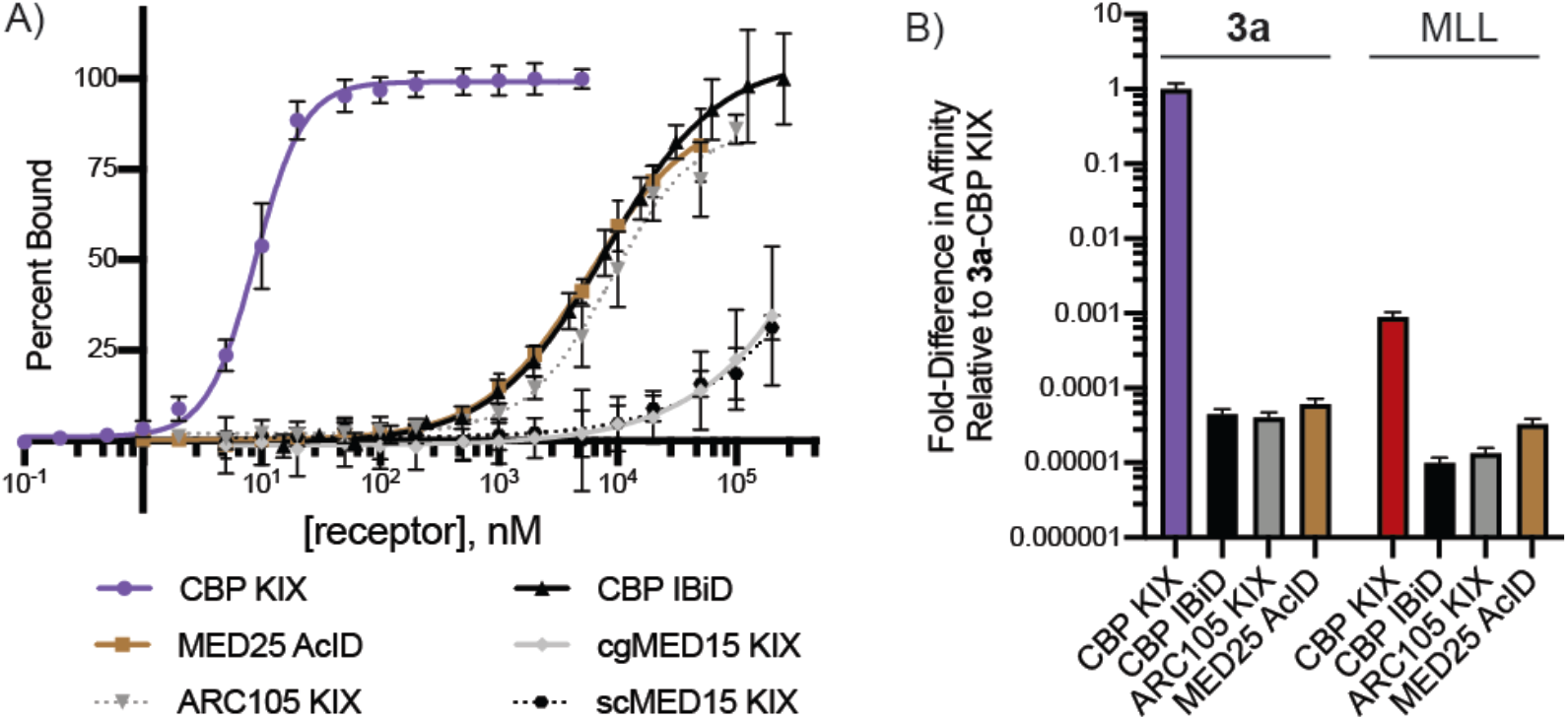
A) Direct-binding fluorescence polarization data of **3a** bound to other coactivator domains. Error is SD calculated from technical triplicates of experimental triplicates. B) Fold difference in dissociation constants for **3a** and MLL relative to **3a**-CBP KIX interaction. Error is SD derived from above polarization data or stopped-flow fluorescence experiments.

The MybLL-tide engages full-length CBP and p300 in the context of the proteome. Affinity pulldowns were performed on lysates of the acute myeloid leukemia cell line, MV4-11. Intact CBP and p300 were both successfully pulled down using NeutrAvidin-bound biotinylated MybLL-tide **5**, while biotinylated mutant **6**, and a blank bead control failed to interact with CBP or p300 (Figure 4A, 4B). Furthermore, affinity pulldown of CBP and p300 with **5** was 84-89% inhibited by the addition of 200 nM free MybLL-tide **3b** to the lysate, but no measurable inhibition was observed when 20 μM MybLL-tide **4b** was added to the lysate (Figure S6).

**Figure 4:**
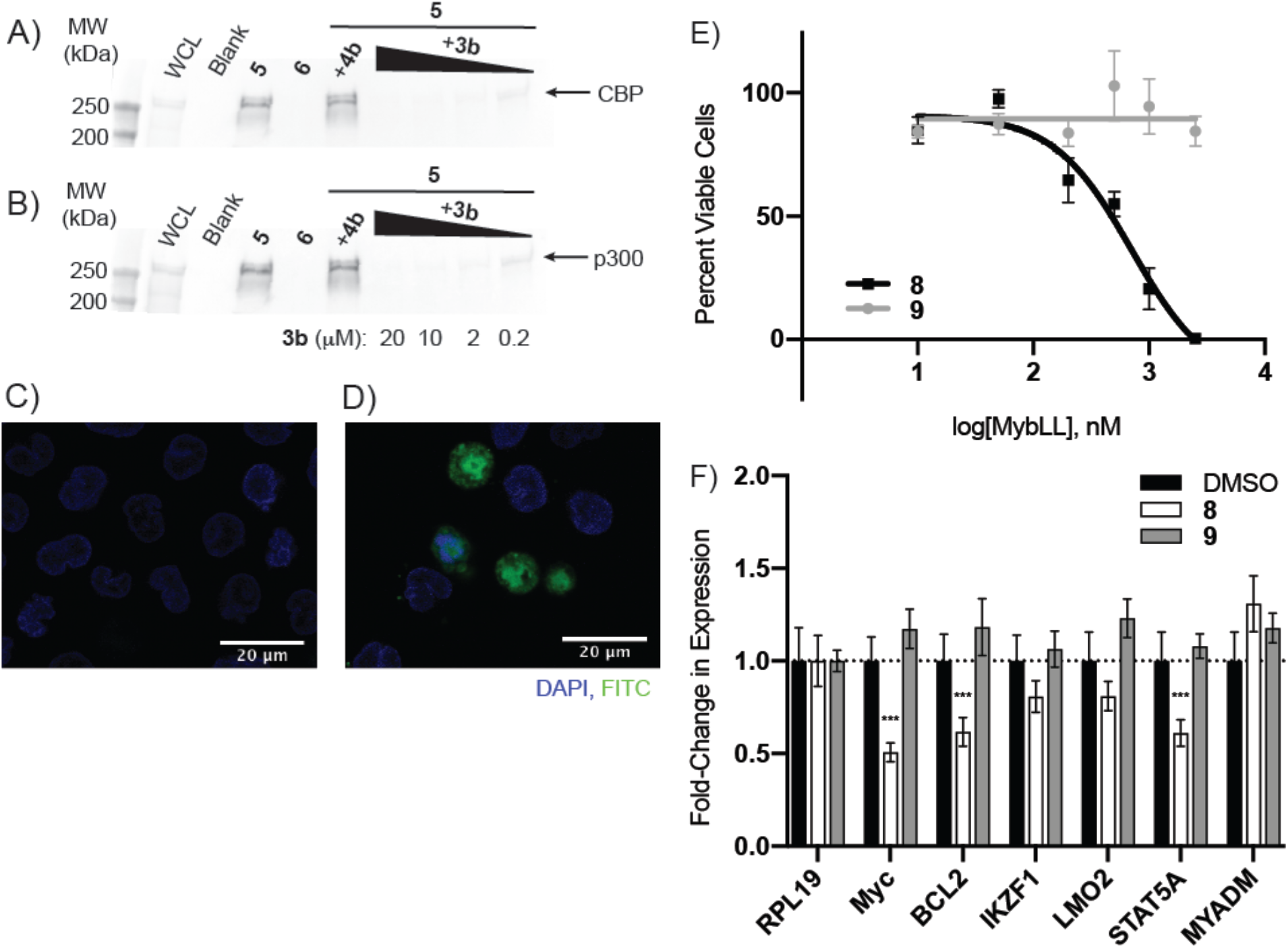
Western blots using antibodies specific for endogenous, full-length A) CBP and B) p300 from MV4-11 whole cell lysates (WCL) showing affinity pulldown of CBP and p300 via biotinylated MybLL-tides **5** and **6** and competitive inhibition of **5** by MybLL-tides **3b** and **4b**. Cell entry of 1 µM FITC-tagged C) MybLL-tide **3a** and D) TAT-MybLL-tide **7** into MV4-11 cells over 6 h. E) Seven-day MTT viability assay on TAT-MybLL-tides **8** and **9** with MV4-11 cells. Error is SD calculated from biological triplicates. F) qRT-PCR analysis of Myb-dependent genes in MV4-11 cells after 6 h exposure to 1 µM TAT-MybLL-tides **8** and **9**. RPL19 is the reference gene. ^***^*p* < 0.001. Error is SD calculated from technical triplicates of biological duplicates.

Cell-active versions of MybLL-tide were developed through incorporation of cell-penetrating peptide (CPP) moieties. A TAT CPP was appended after a short diglycine spacer to the N-terminus of MybLL-tide and the control mutant MybLL-tide (Table 1, compounds **7**-**9**). Initial cell uptake and behavior of TAT-MybLL-tides was assessed using FITC-tagged TAT-MybLL-tide **7** and monitored via confocal microscopy. TAT-MybLL-tide **7** is able to enter cells and nuclei whereas MybLL-tide **3a** shows no measurable cell entry (Fig. 4C, D). As would be expected for a CBP/p300 KIX inhibitor,^18,33,34,38^ TAT-MybLL-tide **8** decreased viability of acute myeloid leukemia cells (MV4-11). Using an MTT assay, **8** inhibited viability of MV4-11 cells over a one-week experiment with an IC_50_ of 700 ± 300 nM (Figure 4E). In contrast, mutant TAT-MybLL-tide **9** did not significantly attenuate viability during the same experiment, nor did **3b** (Figure S8), suggesting negligible off-target toxicity from the TAT CPP or the MybLL-tide architecture itself.

The activity of **8** on MV4-11 cells is also observable over a much shorter time frame when assessing the transcription of Myb-dependent genes. We monitored six genes known to be dependent on Myb activity, but with varying degrees of reported dependence on the Myb•KIX interaction.^50,51^ Of these genes, five (Myc, BCL2, IKZF1, LMO2, and STAT5A) showed moderate to significant down-regulation after 6 hours in the presence of 1 µM **8** (Figure 4F) with Myc, BCL2, and STAT5A experiencing a 1.6 to 2-fold decline; this effect was not observed with the negative control **9** (Figure 4F). This data is consistent with the cell viability data, as 100 nM **8** showed little to no effect on transcription of Myb-dependent genes (Figure S9). Importantly, the sixth gene, MYADM, exhibited little difference between both **8** and **9** suggesting that although it is a Myb target gene, MYADM expression may be less dependent on the Myb•KIX interaction. While further study is being conducted to better glean the differing behavior of these genes, this data suggests that cell penetrating MybLL-tides can be employed to assess Myb-dependent transcriptional changes on a short time scale and can be used to better distinguish cellular processes that are functionally controlled by the CBP/p300 KIX domain from those involving the other domains of the coactivators.

Taken together, these results reported here provide compelling evidence that MybLL-tides potently inhibit the Myb-CBP/p300 interaction, possess exceptional selectivity for the CBP/p300 KIX domain, and are capable of entering cells and interacting with endogenous CBP/p300. Ongoing work is currently focused on improving the potency of cell-active MybLL-tides through optimization of proteolytic stability and cellular entry, while continuing to assess how inhibition of Myb-CBP/p300 alters processes such as cell cycle progression in AML cells. Beyond AML, Myb has also shown aberrant behavior in a number of different cancers, suggesting that MybLL-tide’s ability to abrogate the Myb-CBP/p300 KIX interaction may have broad applications to study a number of different disease states.^19–22^ Future work will focus on deploying MybLL-tide against AML, adenoid cystic carcinoma, and breast cancer to further assess the downstream impacts of inhibiting the Myb-CBP/p300 interaction. Additionally, the overall design strategy may be applicable to the many other coactivators with multiple binding sites and relatively low-affinity binding partners.

## Acknowledgements

The authors acknowledge financial support from the National Institutes of Health (CA242018 to A.K.M.; NIH T32GM008597 to support S.D.S.), and the National Science Foundation (NSF REU (BIO/DBI) # 1560096 to support IAV). We are grateful to Prof. Meghan Breen for the MED15 KIX protein samples, Jean Lodge for ARC105 KIX, Julie Garlick for CBP TAZ1, and Steven Sturlis for MED25 AcID. The authors thank Yejun Liu, Yanira Rodriguez Valdes, and Maya Bulos for their useful input at all stages of the project.

---

Table of contents graphic

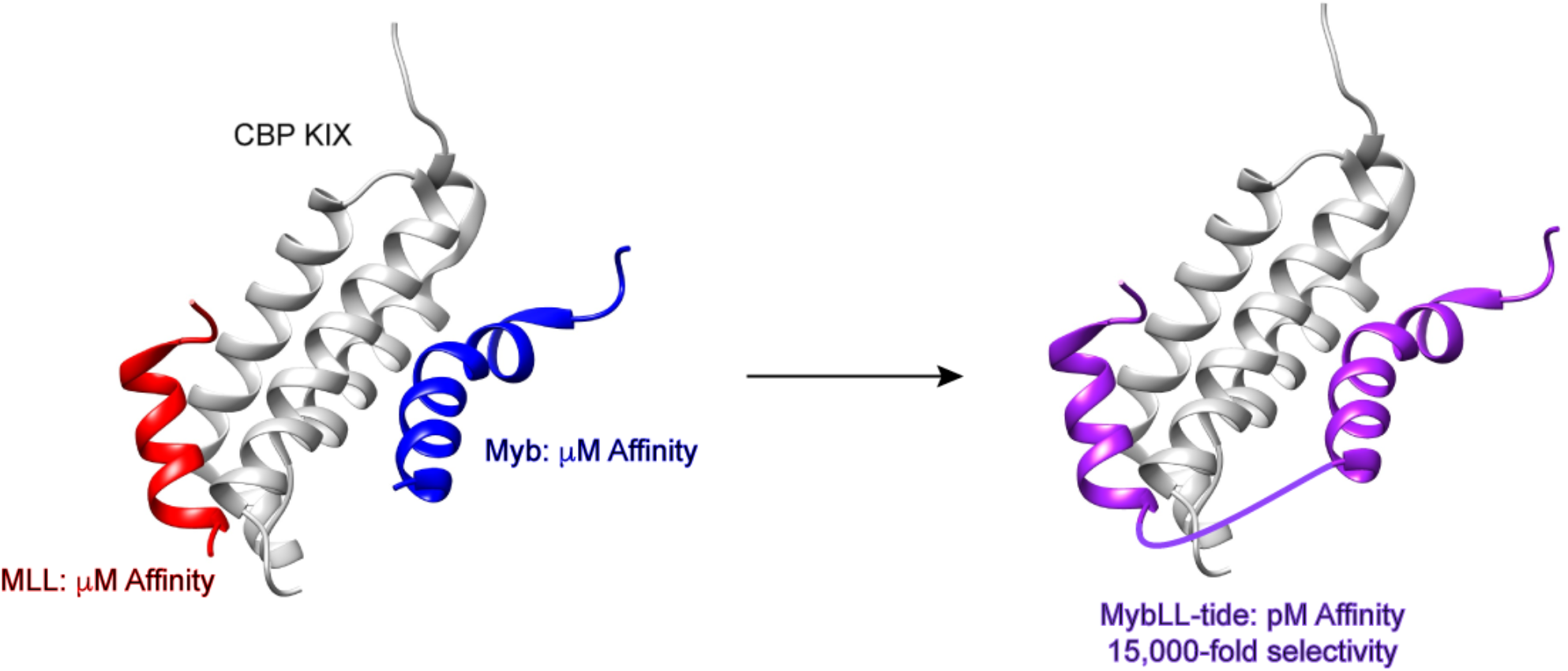

## References

(1) Wang, F.; Marshall, C. B.; Ikura, M. Transcriptional/Epigenetic Regulator CBP/P300 in Tumorigenesis: Structural and Functional Versatility in Target Recognition. Cell. Mol. Life Sci. 2013, 70 (21), 3989–4008. https://doi.org/10.1007/s00018-012-1254-4.

(2) Goodman, R. H.; Smolik, S. CBP/P300 in Cell Growth, Transformation, and Development. Genes Dev. 2000, 14, 1553–1577.

(3) Goto, N. K.; Zor, T.; Martinez-Yamout, M.; Dyson, H. J.; Wright, P. E. Cooperativity in Transcription Factor Binding to the Coactivator CREB-Binding Protein (CBP). J. Biol. Chem. 2002, 277 (45), 43168–43174. https://doi.org/10.1074/jbc.M207660200.

(4) Ogryzko, V. V.; Schiltz, R. L.; Russanova, V.; Howard, B. H.; Nakatani, Y. The Transcriptional Coactivators P300 and CBP Are Histone Acetyltransferases. Cell 1996, 87 (5), 953–959. https://doi.org/10.1016/S0092-8674(00)82001-2.

(5) Breen, M. E.; Mapp, A. K. Modulating the Masters: Chemical Tools to Dissect CBP and P300 Function. Curr. Opin. Chem. Biol. 2018, 45, 195–203. https://doi.org/10.1016/j.cbpa.2018.06.005.

(6) Thakur, J. K.; Yadav, A.; Yadav, G. Molecular Recognition by the KIX Domain and Its Role in Gene Regulation. Nucleic Acids Res. 2014, 42 (4), 2112–2125. https://doi.org/10.1093/nar/gkt1147.

(7) Radhakrishnan, I.; Pérez-Alvarado, G. C.; Parker, D.; Dyson, H. J.; Montminy, M. R.; Wright, P. E. Solution Structure of the KIX Domain of CBP Bound to the Transactivation Domain of CREB: A Model for Activator:Coactivator Interactions. Cell 1997, 91 (6), 741–752. https://doi.org/10.1016/S0092-8674(00)80463-8.

(8) Zor, T.; De Guzman, R. N.; Dyson, H. J.; Wright, P. E. Solution Structure of the KIX Domain of CBP Bound to the Transactivation Domain of C-Myb. J. Mol. Biol. 2004, 337 (3), 521–534. https://doi.org/10.1016/j.jmb.2004.01.038.

(9) Arai, M.; Dyson, H. J.; Wright, P. E. Leu628 of the KIX Domain of CBP Is a Key Residue for the Interaction with the MLL Transactivation Domain. FEBS Lett. 2010, 584 (22), 4500–4504. https://doi.org/10.1016/j.febslet.2010.10.024.

(10) Yang, K.; Stanfield, R. L.; Martinez-Yamout, M. A.; Dyson, H. J.; Wilson, I. A.; Wright, P. E. Structural Basis for Cooperative Regulation of KIX-Mediated Transcription Pathways by the HTLV- 1 HBZ Activation Domain. Proc. Natl. Acad. Sci. 2018, 115 (40), 10040–10045. https://doi.org/10.1073/pnas.1810397115.

(11) Dai, P.; Akimaru, H.; Tanaka, Y.; Hou, D. X.; Yasukawa, T.; Kanei-Ishii, C.; Takahashi, T.; Ishii, S. CBP as a Transcriptional Coactivator of C-Myb. Genes Dev. 1996, 10 (5), 528–540. https://doi.org/10.1101/gad.10.5.528.

(12) Pattabiraman, D. R.; McGirr, C.; Shakhbazov, K.; Barbier, V.; Krishnan, K.; Mukhopadhyay, P.; Hawthorne, P.; Trezise, A.; Ding, J.; Grimmond, S. M.; Papathanasiou, P.; Alexander, W. S.; Perkins, A. C.; Levesque, J.-P.; Winkler, I. G.; Gonda, T. J. Interaction of C-Myb with P300 Is Required for the Induction of Acute Myeloid Leukemia (AML) by Human AML Oncogenes. Blood 2014, 123 (17), 2682–2690. https://doi.org/10.1182/blood-2012-02-413187.

(13) Giotopoulos, G.; Chan, W.-I.; Horton, S. J.; Ruau, D.; Gallipoli, P.; Fowler, A.; Crawley, C.; Papaemmanuil, E.; Campbell, P. J.; Göttgens, B.; Van Deursen, J. M.; Cole, P. A.; Huntly, B. J. P. The Epigenetic Regulators CBP and P300 Facilitate Leukemogenesis and Represent Therapeutic Targets in Acute Myeloid Leukemia. Oncogene 2016, 35 (3), 279–289. https://doi.org/10.1038/onc.2015.92.

(14) Niu, X.; Wang, G.; Wang, Y.; Caldwell, J. T.; Edwards, H.; Xie, C.; Taub, J. W.; Li, C.; Lin, H.; Ge, Y. Acute Myeloid Leukemia Cells Harboring MLL Fusion Genes or with the Acute Promyelocytic Leukemia Phenotype Are Sensitive to the Bcl-2-Selective Inhibitor ABT-199. Leukemia 2014, 28 (7), 1557–1560. https://doi.org/10.1038/leu.2014.72.

(15) Prange, K. H. M.; Mandoli, A.; Kuznetsova, T.; Wang, S.-Y.; Sotoca, A. M.; Marneth, A. E.; van der Reijden, B. A.; Stunnenberg, H. G.; Martens, J. H. A. MLL-AF9 and MLL-AF4 Oncofusion Proteins Bind a Distinct Enhancer Repertoire and Target the RUNX1 Program in 11q23 Acute Myeloid Leukemia. Oncogene 2017, 36 (23), 3346–3356. https://doi.org/10.1038/onc.2016.488.

(16) Somervaille, T. C. P.; Cleary, M. L. Identification and Characterization of Leukemia Stem Cells in Murine MLL-AF9 Acute Myeloid Leukemia. Cancer Cell 2006, 10 (4), 257–268. https://doi.org/10.1016/j.ccr.2006.08.020.

(17) Mitra, P. Transcription Regulation of MYB: A Potential and Novel Therapeutic Target in Cancer. Ann. Transl. Med. 2018, 6 (22), 443–443. https://doi.org/10.21037/atm.2018.09.62.

(18) Uttarkar, S.; Dukare, S.; Bopp, B.; Goblirsch, M.; Jose, J.; Klempnauer, K.-H. Naphthol AS-E Phosphate Inhibits the Activity of the Transcription Factor Myb by Blocking the Interaction with the KIX Domain of the Coactivator P300. Mol. Cancer Ther. 2015, 14 (6), 1276–1285. https://doi.org/10.1158/1535-7163.MCT-14-0662.

(19) Liu, X.; Xu, Y.; Han, L.; Yi, Y. Reassessing the Potential of Myb-Targeted Anti-Cancer Therapy. J. Cancer 2018, 9 (7), 1259–1266. https://doi.org/10.7150/jca.23992.

(20) Costa, A. F.; Altemani, A.; García-Inclán, C.; Fresno, F.; Suárez, C.; Llorente, J. L.; Hermsen, M. Analysis of MYB Oncogene in Transformed Adenoid Cystic Carcinomas Reveals Distinct Pathways of Tumor Progression. Lab. Invest. 2014, 94 (6), 692–702. https://doi.org/10.1038/labinvest.2014.59.

(21) Ho, A. S.; Kannan, K.; Roy, D. M.; Morris, L. G. T.; Ganly, I.; Katabi, N.; Ramaswami, D.; Walsh, A.; Eng, S.; Huse, J. T.; Zhang, J.; Dolgalev, I.; Huberman, K.; Heguy, A.; Viale, A.; Drobnjak, M.; Leversha; M. A., Rice, C. E.; Singh, B.; Iyer, N. G.; Leemans, C. R.; Bloemena, E.; Ferris, R. L.; Seethala, R. R.; Gross, B. E.; Liang, Y.; Sinha, R.; Peng, L.; Raphael, B. J.; Turcan, S.; Gong, Y.; Schultz, N.; Kim, S.; Chiosea, S.; Shah, J. P.; Sander, C.; Lee, W.; Chan, T. A. The Mutational Landscape of Adenoid Cystic Carcinoma. Nat. Genet. 2013, 45 (7), 791–798. https://doi.org/10.1038/ng.2643.

(22) Drabsch, Y.; Ramsay, R. G.; Gonda, T. J. MYB Suppresses Differentiation and Apoptosis of Human Breast Cancer Cells. Breast Cancer Res. 2010, 12 (4), R55. https://doi.org/10.1186/bcr2614.

(23) Ramsay, R. G.; Gonda, T. J. MYB Function in Normal and Cancer Cells. Nat. Rev. Cancer 2008, 8 (7), 523–534. https://doi.org/10.1038/nrc2439.

(24) Li, Y.; Jin, K.; van Pelt, G. W.; van Dam, H.; Yu, X.; Mesker, W. E.; ten Dijke, P.; Zhou, F.; Zhang, L. C-Myb Enhances Breast Cancer Invasion and Metastasis through the Wnt/β-Catenin/Axin2 Pathway. Cancer Res. 2016, 76 (11), 3364–3375. https://doi.org/10.1158/0008-5472.CAN-15-2302.

(25) De Guzman, R. N.; Goto, N. K.; Dyson, H. J.; Wright, P. E. Structural Basis for Cooperative Transcription Factor Binding to the CBP Coactivator. J. Mol. Biol. 2006, 355 (5), 1005–1013. https://doi.org/10.1016/j.jmb.2005.09.059.

(26) Wang, F.; Marshall, C. B.; Yamamoto, K.; Li, G.-Y.; Gasmi-Seabrook, G. M. C.; Okada, H.; Mak, T. W.; Ikura, M. Structures of KIX Domain of CBP in Complex with Two FOXO3a Transactivation Domains Reveal Promiscuity and Plasticity in Coactivator Recruitment. Proc. Natl. Acad. Sci. 2012, 109 (16), 6078–6083. https://doi.org/10.1073/pnas.1119073109.

(27) Lee, C. W.; Arai, M.; Martinez-Yamout, M. A.; Dyson, H. J.; Wright, P. E. Mapping the Interactions of the P53 Transactivation Domain with the KIX Domain of CBP ^†^. Biochemistry 2009, 48 (10), 2115–2124. https://doi.org/10.1021/bi802055v.

(28) Thakur, J. K.; Arthanari, H.; Yang, F.; Pan, S.-J.; Fan, X.; Breger, J.; Frueh, D. P.; Gulshan, K.; Li, D. K.; Mylonakis, E.; Struhl, K.; Moye-Rowley, W. S.; Cormack, B. P.; Wagner, G.; Näär, A. M. A Nuclear Receptor-like Pathway Regulating Multidrug Resistance in Fungi. Nature 2008, 452 (7187), 604–609. https://doi.org/10.1038/nature06836.

(29) Nishikawa, J. L.; Boeszoermenyi, A.; Vale-Silva, L. A.; Torelli, R.; Posteraro, B.; Sohn, Y.-J.; Ji, F.; Gelev, V.; Sanglard, D.; Sanguinetti, M.; Sadreyev, R. I.; Mukherjee, G.; Bhyravabhotla, J.; Buhrlage, S. J.; Gray, N. S.; Wagner, G.; Näär, A. M.; Arthanari, H. Inhibiting Fungal Multidrug Resistance by Disrupting an Activator–Mediator Interaction. Nature 2016, 530 (7591), 485–489. https://doi.org/10.1038/nature16963.

(30) Yang, F.; Vought, B. W.; Satterlee, J. S.; Walker, A. K.; Jim Sun, Z.-Y., Watts, J. L.; DeBeaumont, R.; Mako Saito, R.; Hyberts, S. G.; Yang, S.; Macol, C.; Iyer, L.; Tjian, R.; van den Heuvel, S.; Hart, A. C.; Wagner, G.; Näär, A. M. An ARC/Mediator Subunit Required for SREBP Control of Cholesterol and Lipid Homeostasis. Nature 2006, 442 (7103), 700–704. https://doi.org/10.1038/nature04942.

(31) Jedidi, I.; Zhang, F.; Qiu, H.; Stahl, S. J.; Palmer, I.; Kaufman, J. D.; Nadaud, P. S.; Mukherjee, S.; Wingfield, P. T.; Jaroniec, C. P.; Hinnebusch, A. G. Activator Gcn4 Employs Multiple Segments of Med15/Gal11, Including the KIX Domain, to Recruit Mediator to Target Genes in Vivo. J. Biol. Chem. 2010, 285 (4), 2438–2455. https://doi.org/10.1074/jbc.M109.071589.

(32) Kassube, S. A.; Jinek, M.; Fang, J.; Tsutakawa, S.; Nogales, E. Structural Mimicry in Transcription Regulation of Human RNA Polymerase II by the DNA Helicase RECQL5. Nat. Struct. Mol. Biol. 2013, 20 (7), 892–899. https://doi.org/10.1038/nsmb.2596.

(33) Ramaswamy, K.; Forbes, L.; Minuesa, G.; Gindin, T.; Brown, F.; Kharas, M. G.; Krivtsov, A. V.; Armstrong, S. A.; Still, E.; de Stanchina, E.; Knoechel, B.; Koche, R.; Kentsis, A. Peptidomimetic Blockade of MYB in Acute Myeloid Leukemia. Nat. Commun. 2018, 9 (1), 110. https://doi.org/10.1038/s41467-017-02618-6.

(34) Uttarkar, S.; Dassé, E.; Coulibaly, A.; Steinmann, S.; Jakobs, A.; Schomburg, C.; Trentmann, A.; Jose, J.; Schlenke, P.; Berdel, W. E.; Schmidt, T. J.; Müller-Tidow, C.; Frampton, J.; Klempnauer, K.-H. Targeting Acute Myeloid Leukemia with a Small Molecule Inhibitor of the Myb/P300 Interaction. Blood 2016, 127 (9), 1173–1182. https://doi.org/10.1182/blood-2015-09-668632.

(35) Volkman, H. M.; Rutledge, S. E.; Schepartz, A. Binding Mode and Transcriptional Activation Potential of High Affinity Ligands for the CBP KIX Domain. J. Am. Chem. Soc. 2005, 127 (13), 4649–4658. https://doi.org/10.1021/ja042761y.

(36) Frangioni, J. V.; LaRiccia, L. M.; Cantley, L. C.; Montminy, M. R. Minimal Activators That Bind to the KIX Domain of P300/CBP Identified by Phage Display Screening. Nat. Biotechnol. 2000, 18 (10), 1080–1085. https://doi.org/10.1038/80280.

(37) Takao, S.; Forbes, L.; Uni, M.; Cheng, S.; Pineda, J. M. B.; Tarumoto, Y.; Cifani, P.; Minuesa, G.; Chen, C.; Kharas, M. G.; Bradley, R. K.; Vakoc, C. R.; Koche, R. P.; Kentsis, A. Convergent Organization of Aberrant MYB Complex Controls Oncogenic Gene Expression in Acute Myeloid Leukemia. eLife 2021, 10, e65905. https://doi.org/10.7554/eLife.65905.

(38) Majmudar, C. Y.; Højfeldt, J. W.; Arevang, C. J.; Pomerantz, W. C.; Gagnon, J. K.; Schultz, P. J.; Cesa, L. C.; Doss, C. H.; Rowe, S. P.; Vásquez, V.; Tamayo-Castillo, G.; Cierpicki, T.; Brooks, C. L.; Sherman, D. H.; Mapp, A. K. Sekikaic Acid and Lobaric Acid Target a Dynamic Interface of the Coactivator CBP/P300. Angew. Chem. Int. Ed. 2012, 51 (45), 11258–11262. https://doi.org/10.1002/anie.201206815.

(39) Toto, A.; Giri, R.; Brunori, M.; Gianni, S. The Mechanism of Binding of the KIX Domain to the Mixed Lineage Leukemia Protein and Its Allosteric Role in the Recognition of C-Myb: Mechanism of Binding of the KIX Domain. Protein Sci. 2014, 23 (7), 962–969. https://doi.org/10.1002/pro.2480.

(40) Brüschweiler, S.; Konrat, R.; Tollinger, M. Allosteric Communication in the KIX Domain Proceeds through Dynamic Repacking of the Hydrophobic Core. ACS Chem. Biol. 2013, 8 (7), 1600–1610. https://doi.org/10.1021/cb4002188.

(41) Palazzesi, F.; Barducci, A.; Tollinger, M.; Parrinello, M. The Allosteric Communication Pathways in KIX Domain of CBP. Proc. Natl. Acad. Sci. 2013, 110 (35), 14237–14242. https://doi.org/10.1073/pnas.1313548110.

(42) Pomerantz, W. C.; Wang, N.; Lipinski, A. K.; Wang, R.; Cierpicki, T.; Mapp, A. K. Profiling the Dynamic Interfaces of Fluorinated Transcription Complexes for Ligand Discovery and Characterization. ACS Chem. Biol. 2012, 7 (8), 1345–1350. https://doi.org/10.1021/cb3002733.

(43) Rooklin, D.; Modell, A. E.; Li, H.; Berdan, V.; Arora, P. S.; Zhang, Y. Targeting Unoccupied Surfaces on Protein–Protein Interfaces. J. Am. Chem. Soc. 2017, 139 (44), 15560–15563. https://doi.org/10.1021/jacs.7b05960.

(44) Lecoq, L.; Raiola, L.; Chabot, P. R.; Cyr, N.; Arseneault, G.; Legault, P.; Omichinski, J. G. Structural Characterization of Interactions between Transactivation Domain 1 of the P65 Subunit of NF-ΚB and Transcription Regulatory Factors. Nucleic Acids Res. 2017, 45 (9), 5564–5576. https://doi.org/10.1093/nar/gkx146.

(45) Pattabiraman, D. R.; Sun, J.; Dowhan, D. H.; Ishii, S.; Gonda, T. J. Mutations in Multiple Domains of C-Myb Disrupt Interaction with CBP/P300 and Abrogate Myeloid Transforming Ability. Mol. Cancer Res. 2009, 7 (9), 1477–1486. https://doi.org/10.1158/1541-7786.MCR-09-0070.

(46) Odoux, A.; Jindal, D.; Tamas, T. C.; Lim, B. W. H.; Pollard, D.; Xu, W. Experimental and Molecular Dynamics Studies Showed That CBP KIX Mutation Affects the Stability of CBP:C-Myb Complex. Comput. Biol. Chem. 2016, 62, 47–59. https://doi.org/10.1016/j.compbiolchem.2016.03.004.

(47) Lin, C. H.; Hare, B. J.; Wagner, G.; Harrison, S. C.; Maniatis, T.; Fraenkel, E. A Small Domain of CBP/P300 Binds Diverse Proteins: Solution Structure and Functional Studies. Mol. Cell 2001, 8, 581–590.

(48) De Guzman, R. N.; Wojciak, J. M.; Martinez-Yamout, M. A.; Dyson, H. J.; Wright, P. E. CBP/P300 TAZ1 Domain Forms a Structured Scaffold for Ligand Binding ^†, ‡^. Biochemistry 2005, 44 (2), 490–497. https://doi.org/10.1021/bi048161t.

(49) Bontems, F.; Verger, A.; Dewitte, F.; Lens, Z.; Baert, J.-L.; Ferreira, E.; Launoit, Y. de; Sizun, C.; Guittet, E.; Villeret, V.; Monté, D. NMR Structure of the Human Mediator MED25 ACID Domain. J. Struct. Biol. 2011, 174 (1), 245–251. https://doi.org/10.1016/j.jsb.2010.10.011.

(50) Lorenzo, P. I.; Brendeford, E. M.; Gilfillan, S.; Gavrilov, A. A.; Leedsak, M.; Razin, S. V.; Eskeland, R.; Saether, T.; Gabrielsen, O. S. Identification of C-Myb Target Genes in K562 Cells Reveals a Role for c-Myb as a Master Regulator. Genes Cancer 2011, 2 (8), 805–817. https://doi.org/10.1177/1947601911428224.

(51) Bengtsen, M.; Klepper, K.; Gundersen, S.; Cuervo, I.; Drabløs, F.; Hovig, E.; Sandve, G. K.; Gabrielsen, O. S.; Eskeland, R. C-Myb Binding Sites in Haematopoietic Chromatin Landscapes. PLOS ONE 2015, 10 (7), e0133280. https://doi.org/10.1371/journal.pone.0133280.

