## Supporting Information for "A Dual-Site Inhibitor of CBP/p300 KIX is a Selective and Effective Modulator of Myb"

### Table of Contents

| | |
|---|---|
| **General** | **2** |
| **Methods** | **3** |
| **Supplemental Figures** | **11** |
| **Peptide Sequences and Mass Spectrometry** | **23** |
| **Structures and Analytical Traces of Novel Peptides** | **25** |
| **References** | **44** |

**General**.

Amino acids and coupling reagents were purchased from CEM, Gyros Protein Technologies, and Chem Impex. Other peptide synthesis reagents including trifluoroacetic acid and piperidine were purchased from Sigma Aldrich. Acetonitrile and other solvents were acquired from Fisher Scientific. MV4-11 cells were acquired from ATCC. All primers were acquired from Sigma. All antibodies were purchased from Abcam or Santa Cruz Biotechnology. PowerUp SYBR Green Master Mix for qPCR was purchased from Applied Biosystems and iScript Reverse Transcriptase was acquired from Bio-Rad.

**Abbreviations**.

| | |
|---|---|
| CBP | CREB-Binding Protein |
| KIX | Kinase-inducible domain Interacting domain |
| AML | Acute Myeloid Leukemia |
| ABD | Activator-Binding Domain |
| TAD | Transcriptional Activation Domain |
| IBiD | Interferon-Binding Domain |
| TAZ1 | Transcription Adaptor putative Zinc finger |
| cgMED15 | *Candida glabrata* MED15 |
| scMED15 | *Saccharomyces cerevisiae* MED15 |
| AcID | Activator Interacting Domain |
| CPP | Cell-Penetrating Peptide |
| PEG | polyethylene glycol |
| AEEAc | 8-amino-3,6-dioxaoctanoic acid |
| PrOF | Protein-Observed Fluorine |
| DAPI | 4',6-diamidino-2-phenylindole |
| FITC | fluorescein isothiocyanate |
| qRT-PCR | quantitative Reverse Transcription-Polymerase Chain Reaction |

**Methods**.

**Peptide Synthesis.** All MybLL-tides, MLL and Myb peptides, and the CBP IBiD domain (2063-2111) were synthesized using standard Fmoc solid phase peptide synthesis on a Liberty Blue synthesizer. FITC-tagged peptides were synthesized by reacting resin-bound peptides with FITC (1.5 eq) in 5% DIPEA in DMF for 16 hr. Peptides were cleaved from resin using 90% trifluoroacetic acid (TFA), 5% thioanisole, 3% ethanedithiol, and 2% anisole for 4 hours. Resin was filtered, the solvent evaporated under nitrogen, and the crude peptides precipitated and pelleted from cold ether. Crude peptides were dissolved in 20% acetonitrile in water, frozen and lyophilized. After lyophilization, TAT-containing peptides were subjected to an additional 2 hours in cleavage cocktail followed by a second round of evaporation, ether precipitation, and lyophilization in order to fully remove protecting groups. Dry crude peptides were re-dissolved in 50 mM Tris (pH = 8.0) with 20% acetonitrile and 25 mM DTT and agitated for 1 hour before being purified via HPLC on a C18 column using an Agilent 1260 HPLC and acetonitrile and 0.1% aqueous TFA as running solvents. Pure fractions were lyophilized and frozen until ready for use. MybLL-tides intended for cell studies were subjected to an additional round of HPLC purification with 0.5% aqueous acetic acid and subsequent lyophilization to remove any residual TFA co-salts. Identity and purity were verified by analytical HPLC and mass spectrometry, obtained using an Agilent 6230 LC/TOF or an Agilent 6545 LC/Q-TOF.

**p300 KIX expression plasmid construction**. Using the CBP GACKIX (586-672) plasmid in a pRSETb vector[1] with an N-terminal 6x HIS tag as a template, p300 KIX (566-652) was designed using standard quick change site-directed mutagenesis to mutate the 9 amino acid that differ from

CBP KIX. The primers for the mutants are listed below. Plasmid sequence identity was verified using standard Sanger sequencing methods at the University of Michigan DNA Sequencing Core.

1) Val587 to Ile567
Fwd: ATG GCT AGC GGT ATT CGA AAA GGC TGG
Rev: CCA GCC TTT TCG AAT ACC GCT AGC CAT

2) Gly590 to Gln570
Fwd: GGT ATT CGA AAA CAG TGG CAT GAA CAT
Rev: ATG TTC ATG CCA CTG TTT TCG AAT ACC

3) His594 to Asp574
Fwd: CAG TGG CAT GAA GAC GTG ACT CAG GAC
Rev: GTC CTG AGT CAC GTC TTC ATG CCA CTG

4) Val595 to Ile575
Fwd: TGG CAT GAA GAC ATC ACT CAG GAC CTA
Rev: TAG GTC CTG AGT GAT GTC TTC ATG CCA

5) Ser601 to Asn581
Fwd: CAG GAC CTA CGG AAC CAT CTA GTC CAT
Rev: ATG GAC TAG ATG GTT CCG TAG GTC CTG

6) Lys633 to Arg613
Fwd: GTT GCC TAT GCT AGG AAA GTG GAG GGA
Rev: TCC CTC CAC TTT CCT AGC ATA GGG AAC

7) Ser645 to Asn625 and Asp647 to Ala627
Fwd: GCT AAT AAC AGG GCC GAA TAC TAT CAT
Rev: ATG ATA GTA TTC GGC CCT GTT ATT AGC

8) Ser660 to Thr650
Fwd: GAA GAA AAG CGG AGG ACC CGT TTA TAG AAG
Rev: CTT CTA TAA ACG GGT CCT CCG CTT TTC TTC

Plasmid Sequence:

GNATTCCCTGAATATTTTGTTTACTTTAGAAGGAGATATACAT ATG CGG GGT TCT
CATCAT CAT CAT CAT CAT GGT ATG GCT AGC GGT ATT CGA AAA CAG TGG CAT
GAA GAC ATC ACT CAG GAC CTA CGG AAC CAT CTA GTC CAT AAA CTC GTT CAA
GCC ATC TTC CCA ACT CCA GAC CCT GCA GCT CTG AAA GAT CGC CGC ATG GAG
AAC CTG GTT GCC TAT GCT AGG AAA GTG GAG GGA GAC ATG TAT GAG TCT GCT
AAT AAC AGG GCC GAA TAC TAT CAT TTA TTA GCA GAG AAA ATC TAT AAA ATA
CAA AAA GAA CTA GAA GAA AAG CGG AGG ACC CGT TTA TAG
AAGCTTGATCCGGCTGCTAACAAAGCCCGAAAGGAAGCTGAGTTG

Amino Acid Sequence (mutated residues highlighted):

MRGSHHHHHHGMASGIRKQWHEDITQDLRNHLVHKLVQAIFPTPDPAALKDRRMENL VAYARKVEGDMYESANNRAEYYHLLAEKIYKIQKELEEKRRTRL

**Protein Expression and Purification.** CBP KIX was expressed and purified as previously described,[1] and the methodology modified to express and purify p300 KIX. The His-tagged p300 KIX domain (566-652) transformed into the pRSETb vector was transfected into BL21-AI cells. Cells were grown to $OD_{600}$ = 0.8, then cooled to 20 °C induced using IPTG and 20% arabinose and held at 20 °C for 16 hours. Cells were pelleted, resuspended in lysis buffer (100 mM Tris, 500 mM NaCl, 5 mM imidazole, 10% glycerol, 10 mM β-mercaptoethanol, pH = 7.4) supplemented with Roche protease inhibitor tablets and lysed using a Fisher Scientific Model 500 sonic dismembrator. Lysate was filtered and the filtered solution incubated with Ni-NTA resin (1 mL resin/10 mL lysate) for 1 hr at 4 °C. Resin was spun down, decanted, and washed three times with wash buffer (100 mM Tris, 500 mM NaCl, 5 mM imidazole, 10 mM β-mercaptoethanol, pH = 7.4) for 5 min/wash. Resin was spun down, supernatant removed, and His-tagged p300 KIX eluted from the resin with elution buffer (100 mM Tris, 500 mM NaCl, 200 mM imidazole, 10 mM β-mercaptoethanol, pH = 7.4) for 5 min. Resin was spun down, supernatant removed, and elution repeated 5-7 times total. Fractions containing p300 KIX were combined, dialyzed into 50 mM sodium phosphate pH 7.2 w/ 1 mM DTT and purified further using Source S ion exchange columns and 50 mM sodium phosphate, 1 M NaCl, pH 7.2, 1 mM DTT elution buffer. p300 KIX was evaluated on an Agilent 6545 LC/Q-TOF (mass found = 12060.76) and dialyzed into 10 mM sodium phosphate (pH = 6.8), 100 mM sodium chloride, 10 % glycerol, and 0.01% NP-40 for storage or directly into experimental buffers. Other proteins used in this study were expressed and purified according to previously reported methods.[2–8]

**Fluorescence Polarization.** Polarization experiments were performed using FITC-tagged peptides and iterative dilutions of CBP KIX (180 μM down to 100 pM) in 10 mM sodium phosphate (pH = 6.8), 100 mM sodium chloride, 10 % glycerol, and 0.01% NP-40 with 10 mM DTT.[9] Experiments were performed in triplicate and analyzed using a BMG Pherastar plate reader. Selectivity experiments were performed using the same buffer system with varying concentrations of different proteins (200 pM up to 200 μM). Dissociation constants were calculated using from $EC_{50}$ values using the equation[10]:

$$K_D = ((R_T*(1-F_B)+L_T*F_B)/ F_B) - L_T$$

Where $R_T$ = protein concentration, $F_B$ = fraction of bound FITC-peptide, and $L_T$ = total FITC-peptide concentration.

**Stopped-flow fluorescence.** Stopped-flow fluorescence experiments were performed at 25 °C in KIX storage buffer (10 mM sodium phosphate, 100 mM NaCl, 10 mM DTT, 10% glycerol, 0.01% NP-40, pH 6.8). All concentrations reported are after mixing. The FITC fluorophore was excited at 488 nm, and fluorescence intensity was measured at wavelengths >510 nm using a long-pass filter (Corion). Concentration dependence experiments to determine $k_{on}$ were completed by mixing of a constant concentration of 0.025 μM FITC-labeled MybLL variant with variable concentrations of excess KIX.[11,12] The value of $k_{off}$ was determined via low concentration 1:1 KIX-MybLL-tide association experiments where 5 nM KIX was rapidly mixed with 5 nM FITC-MybLL-tide (**1a**, **2a**, and **3a**). This approach has been previously used to accurately estimate dissociation constants in the pM range[12] and was used because standard out-competition experiments – where pre-complexed KIX•FITC-MybLL is rapidly mixed with a large excess of unlabeled MybLL –

displayed multiphasic kinetics and aggregation behavior. Typically, 10-15 traces were averaged before fitting.

Traces from concentration dependence experiments were fitted using a single exponential equation (first equation below), where *F(t)* is the fluorescence at time *t*, $F_\infty$ is the endpoint fluorescence, $\Delta F$ is the fluorescence amplitude, and $k_{obs}$ is the observed rate constants. The $k_{obs}$ values were plotted as a function of KIX concentration [KIX] and fit to a linear equation (second equation below) to determine the value of $k_{on}$. The value of $k_{off}$ was included in the fits, but due to the low value it was not possible to estimate $k_{off}$ via these concentration dependence experiments.

$$F(t) = F_\infty + \Delta F * \exp(-k_{obs} \times t)$$

$$k_{obs} = k_{on} * [KIX] + k_{off}$$

Traces from low concentration 1:1 mixing experiments were directly fit to the below equation to obtain $k_{off}$, where $F_0$ is the initial fluorescence and $k_{on}$ was restricted to the value obtained from concentration dependence experiments; all other values are the same as previously defined.

$$F(t) = F_0 + \Delta F \left( \frac{(b - z)(1 - exp(z * k_{on} * t))}{2\left(\left(\frac{b - z}{b + z}\right) exp(z * k_{on} * t) - 1\right)} \right)$$

$$b = -\left(\frac{k_{off}}{k_{on}}\right) - 2 * [KIX]$$

$$z = \sqrt{\left(\frac{k_{off}}{k_{on}}\right)^2 + 4 * \left(\frac{k_{off}}{k_{on}}\right) * [KIX]}$$

**Thermal Melts.** CBP KIX was dialyzed into circular dichroism (CD) buffer (10 mM sodium phosphate, 100 mM sodium fluoride, pH = 6.8) and MybLL-tides were also dissolved in CD buffer. Solutions containing 10 μM CBP KIX, 10 μM 1:1 CBP KIX:MybLL-tides, and 10 μM 1:1:1 CBP KIX:MLL:Myb were all prepared and heated from 10 to 90 °C on a JASCO CD Spectrophotometer while monitoring CD signal at 208 and 222 nm. Melting curves were fit to a sigmoidal curve to calculate melting points.

**PrOF NMR.** CBP KIX labeled with 3-fluorotyrosine (3FY) was expressed and purified as previously described and fluorine-19 NMR resonances assigned in accordance with the literature using a Varian 500 MHz NMR spectrometer with a Varian 5 mm PFG OneNMR Probe.[3] MybLL-tide **3b** was titrated in (0, 0.1, 0.175, 0.25, 0.5, 0.75, 1, and 1.25 eq) and the changes in fluorine chemical shifts > 0.15 ppm used to determine the KIX tyrosine residues impacted by MybLL-tide binding.

**Cell Culture.** MV4-11 cells were grown in 10% fetal bovine serum in Iscove's modified Dulbecco's medium (IMDM) at 37 °C in 5% carbon dioxide.

**Affinity Pulldown.** Neutravidin slurry (Pierce 29200, 100 μL) was centrifuged at 3000 g for 2 min, washed twice with binding buffer (100 mM sodium phosphate, 150 mM sodium chloride, pH = 7.2) and resuspended in binding buffer with 10 nanomoles biotinylated MybLL-tide **5** or **6** and 10 mM DTT. Resin was agitated for 1 h at RT, centrifuged, aspirated, and Superblock added. Resin was agitated for 1 h at RT, centrifuged, aspirated, and washed three times with lysis buffer

(10 mM Tris (pH = 7.4), 150 mM sodium chloride, 0.5 mM EDTA, 1 mM DTT, 10% glycerol, and 0.5% Triton X-100). Meanwhile, MV4-11 cells (10 million cells/pulldown) were washed with cold PBS, pelleted, and aspirated three times. Cells were resuspended in cold lysis buffer (1 mL per 10 million cells) supplemented with protease inhibitors (0.5 mM AEBSF, 0.01 mM Bestatin, 0.1 mM Leupeptin, and 0.001 mM Pepstatin), vortexed vigorously for 10 seconds, incubated on ice for 10 min, then centrifuged at 3000 g for 10 min. Lysate was transferred to microcentrifuge tubes and debris was pelleted via centrifugation at 15,000 g for 10 min. Whole cell lysate was removed and added to resin-bound MybLL-tides with DMSO as a control or inhibitory biotin-free MybLL-tides **3b** (20, 10, 2, and 0.2 μM) or **4b** (20 μM). Resin was agitated for 4 h, washed twice, and the resin-bound components removed through boiling in 5:2:13 Laemli buffer: β-mercaptoethanol: 0.1 M Citric acid (pH = 2.1) before separation on 4-20% PAGE bis-tris gel. The gel was transferred to a PVDF membrane using a Trans-Blot Turbo Transfer System, blocked in Superblock, and blotted using anti-CBP (1:250 sc-7300 from and 1:1000 anti-mouse Igg secondary sc-516102 from Santa Cruz). Blot was stripped using ReStore stripping buffer, blocked, and then blotted with anti-p300 (abcam10485 from Abcam and anti-Rabbit Igg secondary sc-2357 from Santa Cruz). Biotinylated peptides **5** and **6** carried C841S mutations to prevent disulfide formation during resin-binding and in subsequent pulldowns.

**Confocal Microscopy.** MV4-11 cells were counted and resuspended in serum-free media and 500 μL of a $6*10^5$ cells/mL suspension transferred to a 24-well plate. Cells were incubated 2 hr at 37 C. Peptides in DMSO were added and cells resuspended by adding 500 μL 5% FBS in IMDM (final concentration of 1 μM peptide). Cells were incubated for 6 hours at 37 C in 5% carbon dioxide, then transferred to microcentrifuge tubes, spun down, and washed with cold PBS (2x). Cells were fixed in paraformaldehyde and nuclei stained with DAPI before being washed and Zstacks imaged using a NIKON A1 confocal microscope. Images were analyzed using ImageJ. Nuclear localization was determined using DAPI-stained nuclei to create three-dimensional nuclear constructs; overlapping nuclear constructs with green fluorescence channel was used to assess nuclear entry. For this experiment, a C841S mutation was installed for peptides **3a** and **7** to avoid issues with fluorescence quenching.

**Viability Assay**. MV4-11 cells were passaged, resuspended in serum-free IMDM at 50,000 cells/mL, and the suspension added (99 μL) to clear 96-well round bottom plates. Cells were incubated for 1 hr at 37 °C in 5% carbon dioxide. MybLL-tides in DMSO were added (1 μL of 200X stocks) at desired concentrations and each well mixed through addition of 5% FBS in IMDM (100 μL). Cells were incubated for 24 h at 37 °C in 5% carbon dioxide. At 24 h, plates were centrifuged, and aspirated carefully to maintain the cell pellet. Cells were resuspended in serum-free media, supplemented with fresh peptide solution, and mixed with 5% FBS in IMDM as described above. Fresh media and peptides were added every 24 hours for a week. At 178 hr, cells were centrifuged, aspirated, and resuspended in 110 μL of 10:1 2.5% FBS in IMDM: MTT solution. Cells were incubated for 4 h at 37 °C in 5% carbon dioxide before being lysed with 100 μL 10% SDS. Lysates were incubated overnight at 37 °C and absorbance read at 570 nm. Experiments were performed in triplicate.

**Myb-Dependent Gene qRT-PCR.** MV4-11 cells were passaged, resuspended in serum-free IMDM at 200,000 cells/mL, and the suspension added (500 μL) to clear 24-well plates. Suspensions were dosed with 5 μL 200X peptide in DMSO (or DMSO as a control) and the suspension mixed through addition of 5% FBS in IMDM (495 μL). Cells were grown for 6 h at

37 °C in 5% carbon dioxide. Cell suspensions were centrifuged, aspirated, and the mRNA extracted. Extracted mRNA was converted to cDNA before being subjected to qPCR on an Applied Biosystems StepOnePlus Real Time PCR System using PowerUp SYBR Green Master Mix and primers listed in Table S10. All data points are derived from technical triplicates of biological duplicates.

| 2AGH Chain | Distance (Å) |
|---|---|
| 1 | 19.4 |
| 2 | 24.8 |
| 3 | 22 |
| 4 | 23.1 |
| 5 | 22.3 |
| 6 | 18.1 |
| 7 | 21.8 |
| 8 | 20 |
| 9 | 20.5 |
| 10 | 23.2 |
| 11 | 23.7 |
| 12 | 18.1 |
| 13 | 22 |
| 14 | 20.6 |
| 15 | 22.7 |
| 16 | 21.3 |
| 17 | 24.7 |
| 18 | 23.9 |
| 19 | 20.4 |
| 20 | 20.4 |
| Avg | 21.7 |
| Standard Deviation | 1.97 |
| Min | 18.1 |
| Max | 24.8 |

**Table S1**. Distances between the Cα atoms of the K291 Myb residue and the P858 residue of MLL using NMR structure from PDB 2AGH.[13]

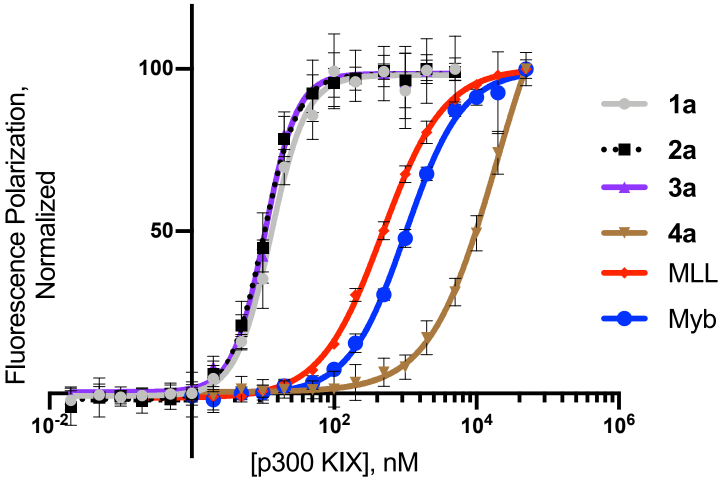


**Figure S1**. Direct-binding fluorescence polarization data of FITC-MybLL-tides, FITC-MLL, and FITC-Myb bound to p300 KIX. Mean ± SD derived from technical triplicates of three independent experiments.

| Peptide | CBP $K_D$ (nM) | p300 $K_D$ (nM) |
|---|---|---|
| **1a** | < 10 | < 10 |
| **2a** | < 10 | < 10 |
| **3a** | < 10 | < 10 |
| **4a** | 16900 ± 1440 | 19000 ± 3520 |
| MLL | 394 ± 20 | 466 ± 10 |
| Myb | 2380 ± 160 | 1000 ± 40 |

**Table S2**. Dissociation constants for FITC-tagged MybLL-tides, MLL, and Myb peptides based on direct-binding fluorescence polarization data.

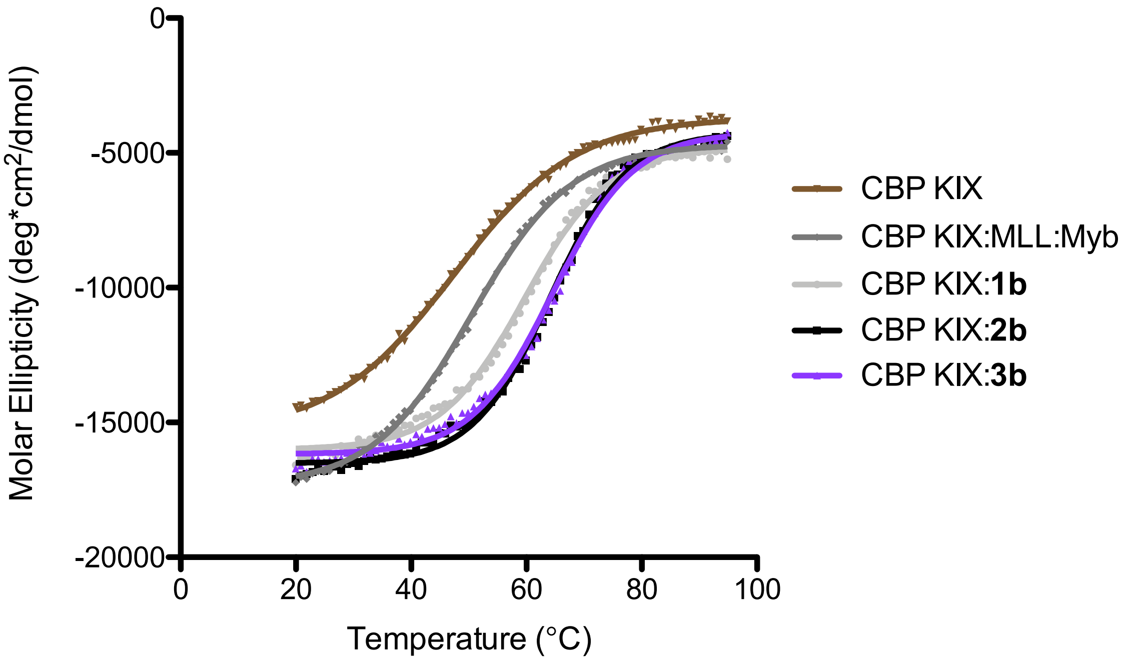


**Figure S2**. Thermal stability of CBP KIX both unbound and in complex with MybLL-tides (**1b**, **2b**, and **3b**), as well as MLL and Myb (ternary complex) as measured by monitoring CD at 222 nm.

| Complex | $T_m$ (ºC) | $\Delta T_m$ (ºC) relative to apo CBP KIX | $\Delta T_m$ (ºC) relative to CBP KIX:MLL:Myb |
|---|---|---|---|
| CBP KIX | 47.3 ± 0.5 | N/A | N/A |
| CBP KIX:MLL:Myb | 50.5 ± 0.3 | 3.2 ± 0.6 | N/A |
| CBP KIX:**1b** | 59.7 ± 0.4 | 12.4 ± 0.6 | 9.2 ± 0.5 |
| CBP KIX:**2b** | 64.1 ± 0.4 | 16.8 ± 0.6 | 13.6 ± 0.5 |
| CBP KIX:**3b** | 64.6 ± 0.5 | 17.3 ± 0.7 | 14.1 ± 0.6 |

**Table S3**. Thermal stability of CBP KIX both unbound and in complex with MybLL-tides (**1b**, **2b**, and **3b**), as well as MLL and Myb (ternary complex).

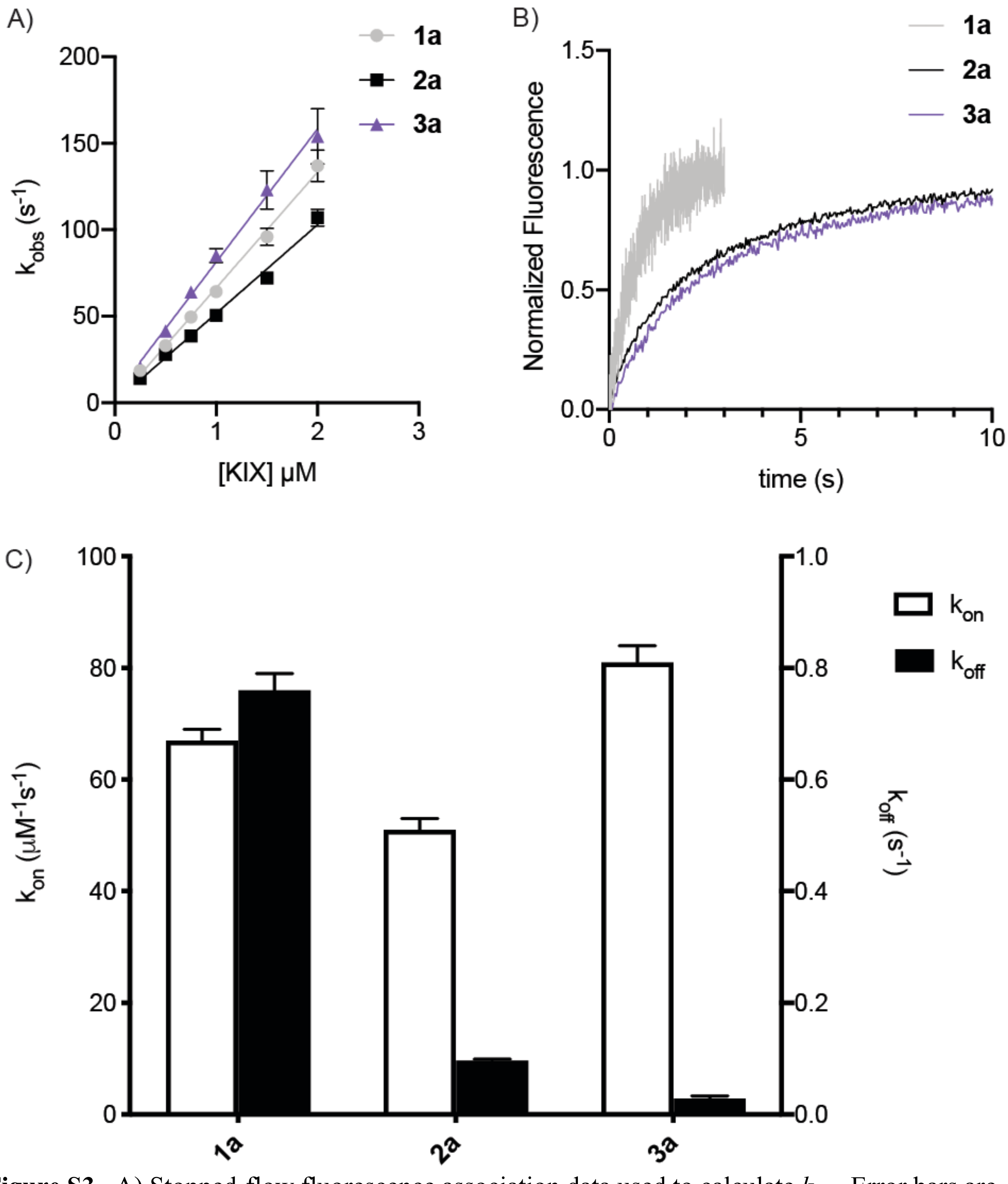


**Figure S3**. A) Stopped-flow fluorescence association data used to calculate $k_{on}$. Error bars are standard deviation derived from three experimental replicates. B) 1:1 mixing experiments of 5 nM FITC-MybLL-tides with 5 nM CBP-KIX used to calculate $k_{off}$. Final calculation was average of 10-15 1:1 association experiments. C) Stopped-flow fluorescence $k_{on}$ and $k_{off}$ for MybLL-tides **1a**, **2a**, and **3a** binding to CBP KIX.

| Peptide | $k_{on}$ ($\mu M^{-1}s^{-1}$) | $k_{off}$ ($s^{-1}$) | $K_D$ (nM) |
|---|---|---|---|
| **1a** | 67 ± 2 | 0.76 ± 0.03 | 11.3 ± 0.6 |
| **2a** | 51 ± 2 | 0.097 ± 0.002 | 1.9 ± 0.08 |
| **3a** | 81 ± 3 | 0.029 ± 0.004 | 0.36 ± 0.05 |

**Table S4**. Rate constants and dissociation constants for MybLL-tides **1a**, **2a**, and **3a** based on stopped-flow fluorescence ($K_D = k_{off}/k_{on}$). Error is standard deviation.

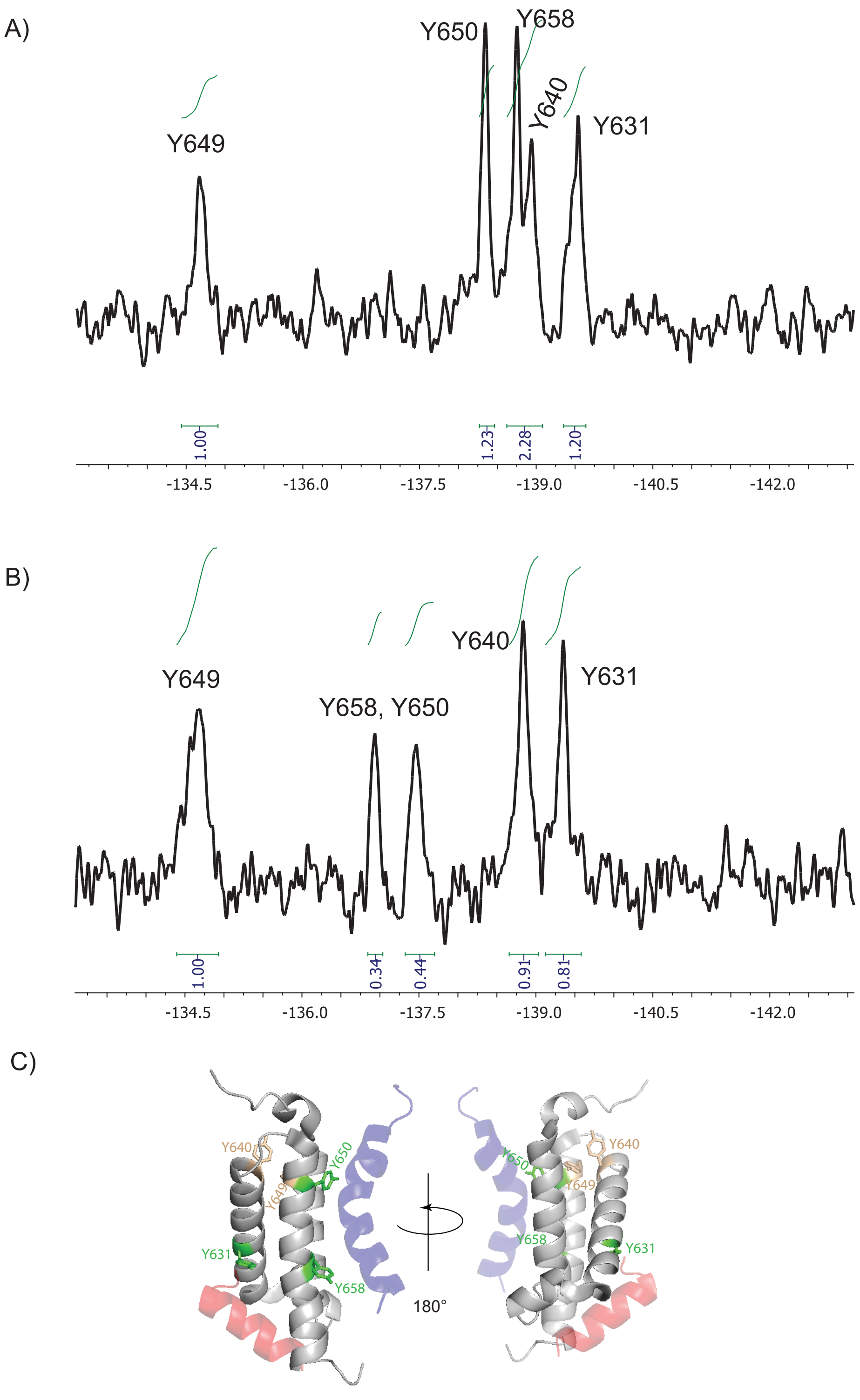


**Figure S4**. Fluorine NMR spectra of A) free 3FY CBP KIX and B) 3FY CBP KIX in complex with MybLL-tide **3b** (1.25 eq). C) Reverse angles of KIX showing the Myb (blue) and MLL (red) binding sites and the tyrosines on each face that are shifting (green) and that are buried and stationary (tan).

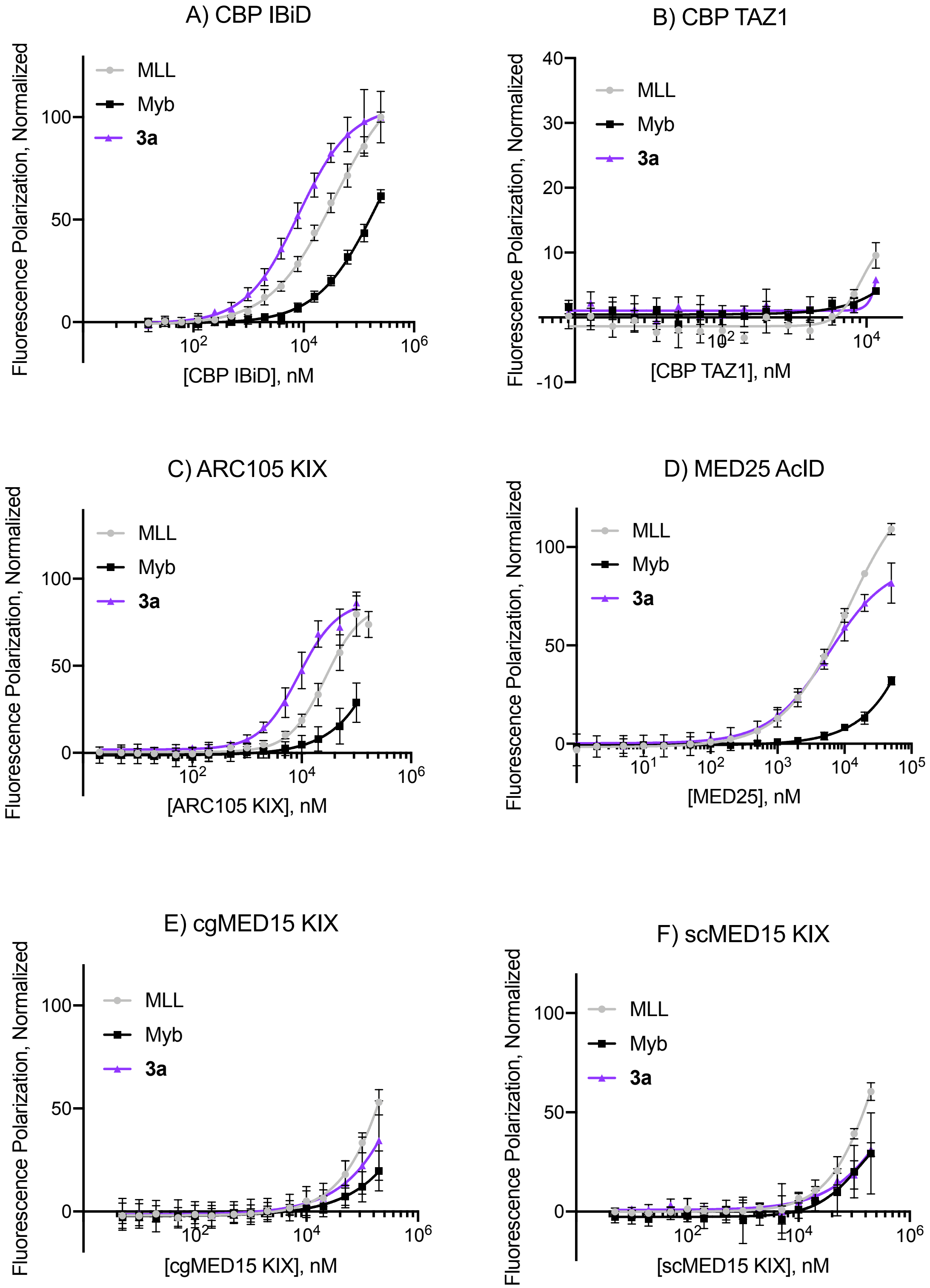


**Figure S5**. Direct-binding fluorescence polarization data for MybLL-tide **3a**, MLL, and Myb binding to A) CBP IBiD, B) CBP TAZ1, C) ARC105 KIX, D) MED25 AcID, E) cgMED15 KIX, and F) scMED15 KIX. Error bars are SD calculated from technical triplicates of three independent experiments, except for CBP TAZ1 which consists of technical triplicates.

| Protein Domain | FITC-MLL (nM) | FITC-Myb (nM) | MybLL-tide **3a** (nM) |
|---|---|---|---|
| CBP KIX | 394 ± 20 | 2380 ± 160 | 0.360 ± 0.05* |
| CBP TAZ1 | > 10000 | > 10000 | > 10000 |
| CBP IBiD | 36100 ± 3800 | > 100000 | 8000 ± 670 |
| ARC105 KIX | 26900 ± 2900 | > 100000 | 8800 ± 700 |
| cgMED15 KIX | > 200000 | > 200000 | > 200000 |
| scMED15 KIX | > 100000 | > 100000 | > 100000 |
| MED25 AcID | 11000 ± 1500 | > 50000 | 5800 ± 530 |

**Table S5**. Dissociation constants for FITC-MLL, FITC-Myb, and FITC-MybLL-tide **3a** with various proteins. All data is derived from above direct-binding fluorescence polarization experiments except: *denotes data for CBP interaction derived from above stopped-flow fluorescence experiments. Mean ± SD.

| Protein Domain | Fold-Change in MLL Binding | Fold-Change in **3a** Binding |
|---|---|---|
| CBP KIX | 1 ± 0.12 | 1120 ± 180 |
| CBP IBiD | 1 ± 0.15 | 4.51 ± 0.61 |
| ARC105 KIX | 1 ± 0.15 | 3.05 ± 0.41 |
| MED25 AcID | 1 ± 0.19 | 1.88 ± 0.30 |

**Table S6**. Fold-change of **3a** binding relative to MLL for CBP KIX, CBP IBiD, ARC105 KIX, and MED25 AcID relative to MLL. Mean ± SD.

| Protein Domain | Fold-Change in Myb Binding | Fold-Change in **3a** Binding |
|---|---|---|
| CBP KIX | 1 ± 0.10 | 6650 ± 1030 |

**Table S7**. Fold-change of **3a** binding relative to Myb for CBP KIX. Mean ± SD.

| Protein Domain | Relative Affinity |
|---|---|
| CBP KIX | 16200 ± 2700 |
| CBP IBiD | 0.73 ± 0.09 |
| ARC105 KIX | 0.66 ± 0.08 |
| MED25 AcID | 1 ± 0.13 |

**Table S8**. Dissociation constants for **3a** normalized to MED25 AcID $K_D$ (nM). Relative Affinity = (MED25 AcID $K_D$)/(Domain $K_D$). Mean ± SD.

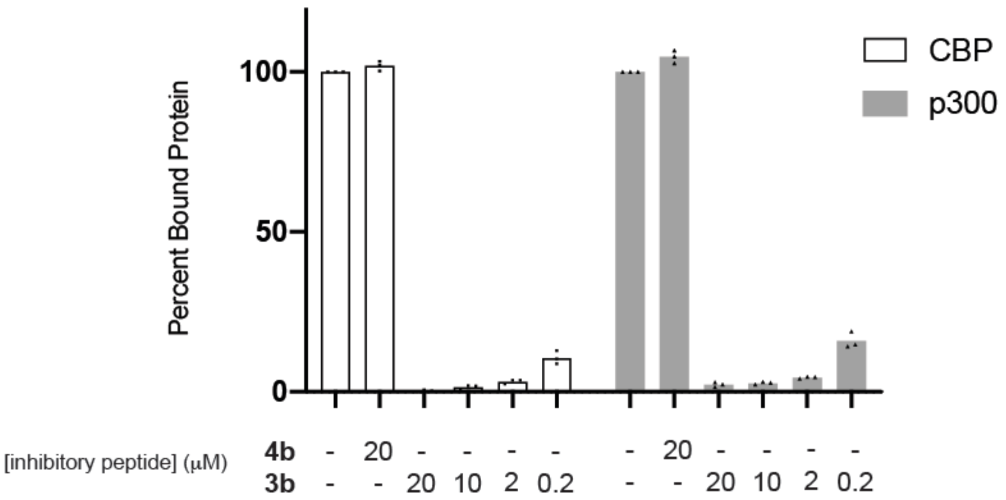


**Figure S6**. Relative quantification of pulldown inhibition as calculated by densitometry. Data is derived from three different exposure times for the blots in figure 2D and E.

| Protein | DMSO | **4b** (20 μM) | **3b** (20 μM) | **3b** (10 μM) | **3b** (2 μM) | **3b** (0.2 μM) |
|---|---|---|---|---|---|---|
| CBP | 100 | 102 ± 2 | 0.3 ± 0.3 | 1.4 ± 0.6 | 3.1 ± 0.6 | 11 ± 2 |
| p300 | 100 | 105 ± 2 | 2.2 ± 0.8 | 2.7 ± 0.4 | 4.4 ± 0.3 | 16 ± 3 |

**Table S9**. Quantification of percentage data in Figure S6.

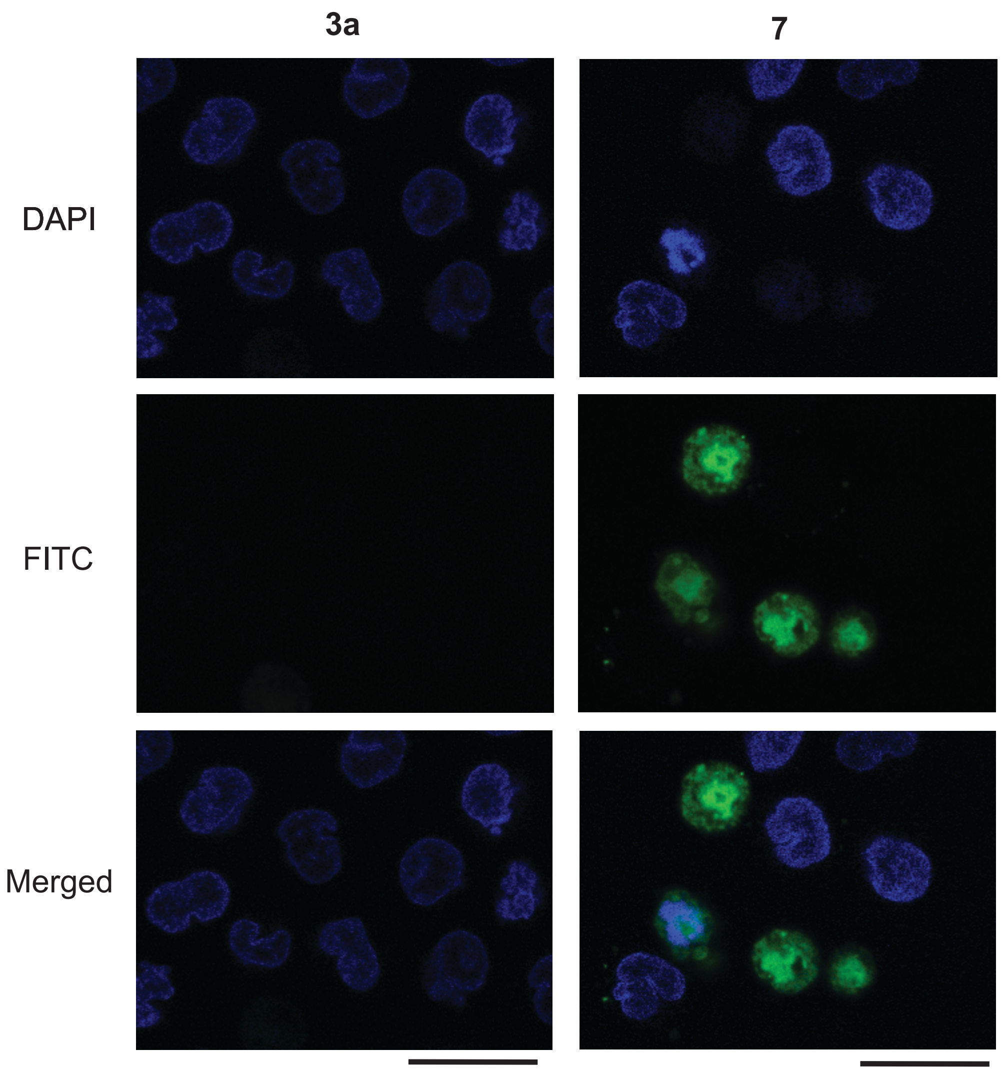


**Figure S7**. Confocal microscopy of MV4-11 cells assessing uptake of 1 μM FITC-MybLL-tides **3a** and **7** (green). Nuclei stained with 4′,6-diamidino-2-phenylindole (DAPI, blue). Scale bar = 20 μm.

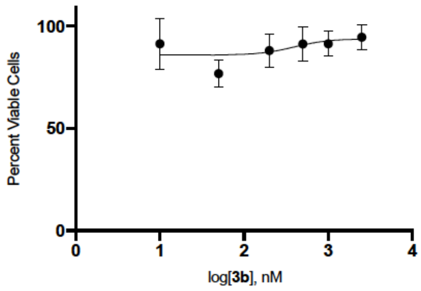


**Figure S8**. MV4-11 cell viability data for **3b**. Mean ± SD from biological triplicates.

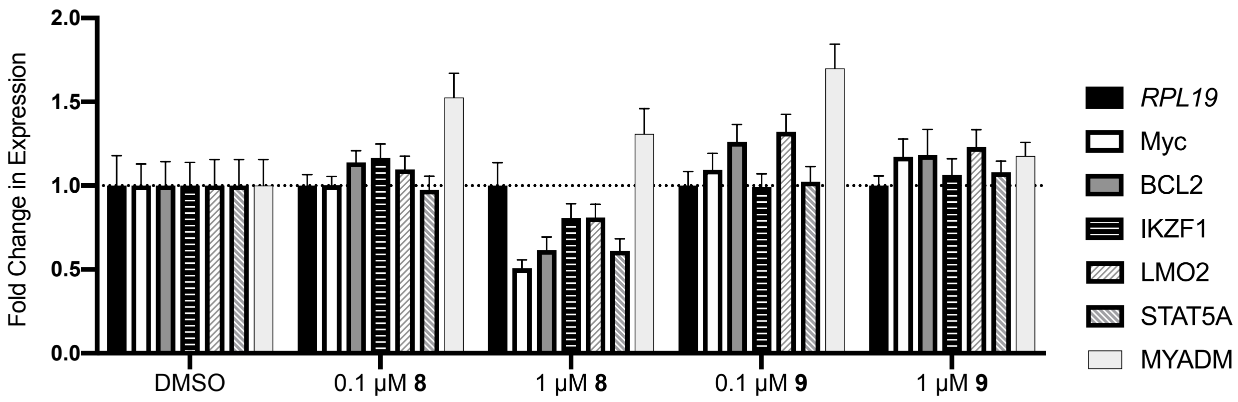


**Figure S9**. MV4-11 6 h qRT-PCR data for **8** and **9** at 0.1 and 1 µM. Error is SD derived from technical triplicates of biological duplicates.

| Primer | Sequence (5'-sequence-3') |
|---|---|
| RPL19 Forward | ATGTATCACAGCCTGTACCTG |
| RPL19 Reverse | TTCTTGGTCTCTTCCTCCTTG |
| Myc Forward | TTCCCCTACCCTCTCAACGACAG |
| Myc Reverse | CCTCATCTTCTTGTTCCTCCTCAG |
| BCL2 Forward | CTGCACCTGACGCCCTTCACC |
| BCL2 Reverse | CACATGACCCCACCGAACTCAAAGA |
| IKZF1 Forward | GCTGCCACAACTACTTGGAAAGC |
| IKZF1 Reverse | AGTCTGTCCAGCACGAGAGATC |
| LMO2 Forward | GCGCCTCTACTACAAACTGGGC |
| LMO2 Reverse | CTCATAGGCACGAATCCGCTTG |
| STAT5A Forward | GTTCAGTGTTGGCAGCAATGAGC |
| STAT5A Reverse | AGCACAGTAGCCGTGGCATTGT |
| MYADM Forward | CCAGTTCGATGAGAAGTATGGCG |
| MYADM Reverse | CAGCCACATACGCCAGTAGGTT |

**Table S10**. Primers used in this study for qRT-PCR.

**Peptide Sequences and Mass Spectrometry**

| Peptide | Peptide Sequence | Expected MW | Observed $[M+4H]^{4+}/4$ | Calculated MW |
|---|---|---|---|---|
| **1a** | FITC-(β-Ala)-DCGNILPSDIMDFVLKNTP-(β-Ala)(AEEAc)$_1$(β-Ala)-KEKRIKELELLLMSTENELKGQQALW-$NH_2$ | 5949.88 | 1488.246 | 5948.95 |
| **1b** | Ac-DCGNILPSDIMDFVLKNTP-(β-Ala)(AEEAc)$_1$(β-Ala)-KEKRIKELELLLMSTENELKGQQALW-$NH_2$ | 5531.45 | 1383.7296 | 5530.89 |
| **2a** | FITC-(β-Ala)-DCGNILPSDIMDFVLKNTP-(β-Ala)(AEEAc)$_2$(β-Ala)-KEKRIKELELLLMSTENELKGQQALW-$NH_2$ | 6095.03 | 1524.7629 | 6094.03 |
| **2b** | Ac-DCGNILPSDIMDFVLKNTP-(β-Ala)(AEEAc)$_2$(β-Ala)-KEKRIKELELLLMSTENELKGQQALW-$NH_2$ | 5676.61 | 1419.9994 | 5676.00 |
| **3a** | FITC-(β-Ala)-DCGNILPSDIMDFVLKNTP-(β-Ala)(AEEAc)$_3$(β-Ala)-KEKRIKELELLLMSTENELKGQQALW-$NH_2$ | 6240.19 | 1561.0295 | 6240.08 |
| **3a** (C841S) | FITC-(β-Ala)-DSGNILPSDIMDFVLKNTP-(β-Ala)(AEEAc)$_3$(β-Ala)-KEKRIKELELLLMSTENELKGQQALW-$NH_2$ | 6224.13 | 1557.0365 | 6224.11 |
| **3b** | Ac-DCGNILPSDIMDFVLKNTP-(β-Ala)(AEEAc)$_3$(β-Ala)-KEKRIKELELLLMSTENELKGQQALW-$NH_2$ | 5821.77 | 1456.2705 | 5821.05 |
| **4a** | FITC-(β-Ala)-DCGNILPSDIMDAVLKNTP-(β-Ala)(AEEAc)$_3$(β-Ala)-KEKRIKELELLAMSTENELKGQQALW-$NH_2$ | 6122.01 | 1531.5162 | 6122.01 |
| **4b** | Ac-DCGNILPSDIMDAVLKNTP-(β-Ala)(AEEAc)$_3$(β-Ala)-KEKRIKELELLAMSTENELKGQQALW-$NH_2$ | 5703.59 | 1426.7618 | 5703.01 |
| **5** | Biotin-(AEEAc)-DSGNILPSDIMDFVLKNTP-(β-Ala)(AEEAc)$_3$(β-Ala)-KEKRIKELELLLMSTENELKGQQALW-$NH_2$ | 6135.12 | 1534.5579 | 6134.20 |
| **6** | Biotin-(AEEAc)-DSGNILPSDIMDAVLKNTP-(β-Ala)(AEEAc)$_3$(β-Ala)-KEKRIKELELLAMSTENELKGQQALW-$NH_2$ | 6016.94 | 1505.0382 | 6016.12 |

**Table S11**. MybLL-tide sequences, expected molecular weight based on structure, observed molecular weight ($[M+4H]^{4+}/4$) on an Agilent LC/TOF, and the calculated molecular weight based on the observed ion. Calculated molecular weight was obtained via deconvolution on an Agilent 1260 TOF.

| Peptide | Peptide Sequence | Expected MW | Observed $[M+8H]^{8+}/8$ | Calculated MW |
|---|---|---|---|---|
| **7** | FITC-(β-Ala)-GRKKRRQRRRPQGGDSGNILPSDIMDFVLKNTP-(β-Ala)(AEEAc)$_3$(β-Ala)-KEKRIKELELLLMSTENELKGQQALW-$NH_2$ | 7958.21 | 995.7718 | 7958.11 |
| **8** | Ac-GRKKRRQRRRPQGGDCGNILPSDIMDFVLKNTP-(β-Ala)(AEEAc)$_3$(β-Ala)-KEKRIKELELLLMSTENELKGQ QALW-$NH_2$ | 7539.79 | 943.5170 | 7540.06 |
| **9** | Ac-GRKKRRQRRRPQGGDCGNILPSDIMDAVLKNTP-(β-Ala)(AEEAc)$_3$(β-Ala)-KEKRIKELELLAMSTENELKGQQALW-$NH_2$ | 7421.61 | 928.6298 | 7420.97 |

**Table S12**. TAT-MybLL-tide sequences, expected molecular weight based on structure, observed molecular weight ($[M+8H]^{8+}/8$) on an Agilent LC/TOF, and the calculated molecular weight based on the observed ion. Calculated molecular weight was obtained using deconvolution on an Agilent 1260 TOF.

| Peptide | Peptide Sequence | Expected MW | Observed $[M+2H]^{2+}/2$ | Calculated MW |
|---|---|---|---|---|
| FITC-MLL | FITC-(β-Ala)-DCGNILPSDIMDFVLKNTP-$NH_2$ | 2551.89 | 1276.5567 | 2551.10 |
| MLL | Ac-DCGNILPSDIMDFVLKNTPY-$NH_2$ | 2296.64 | 1149.0579 | 2296.10 |
| FITC-Myb | FITC-(β-Ala)-KEKRIKELELLLMSTENELKGQQALW-$NH_2$ | 3588.16 | 1794.9788 | 3587.94 |
| Myb | Ac-KEKRIKELELLLMSTENELKGQQALW-$NH_2$ | 3169.74 | 1585.1462 | 3168.76 |
| CBP IBiD (2063-2111) | Ac-SPSALQDLLRTLKSPSSPQQQQQVLNILKSNPQLMAAFIKQRTAKYVAN-$NH_2$ | 5493.39 | 1374.2633† | 5493.01 |

**Table S13**. Myb, MLL, and IBiD peptides, expected molecular weight based on structure, observed molecular weight ($[M+2H]^{2+}/2$) on an Agilent LC/TOF, and the calculated molecular weight based on the observed ion. Calculated molecular weight was obtained using deconvolution on an Agilent 1260 TOF. †Denotes ($[M+4H]^{4+}/4$) ion observed for CBP IBiD.

### Structures and Analytical Traces of Peptides Used in this Study

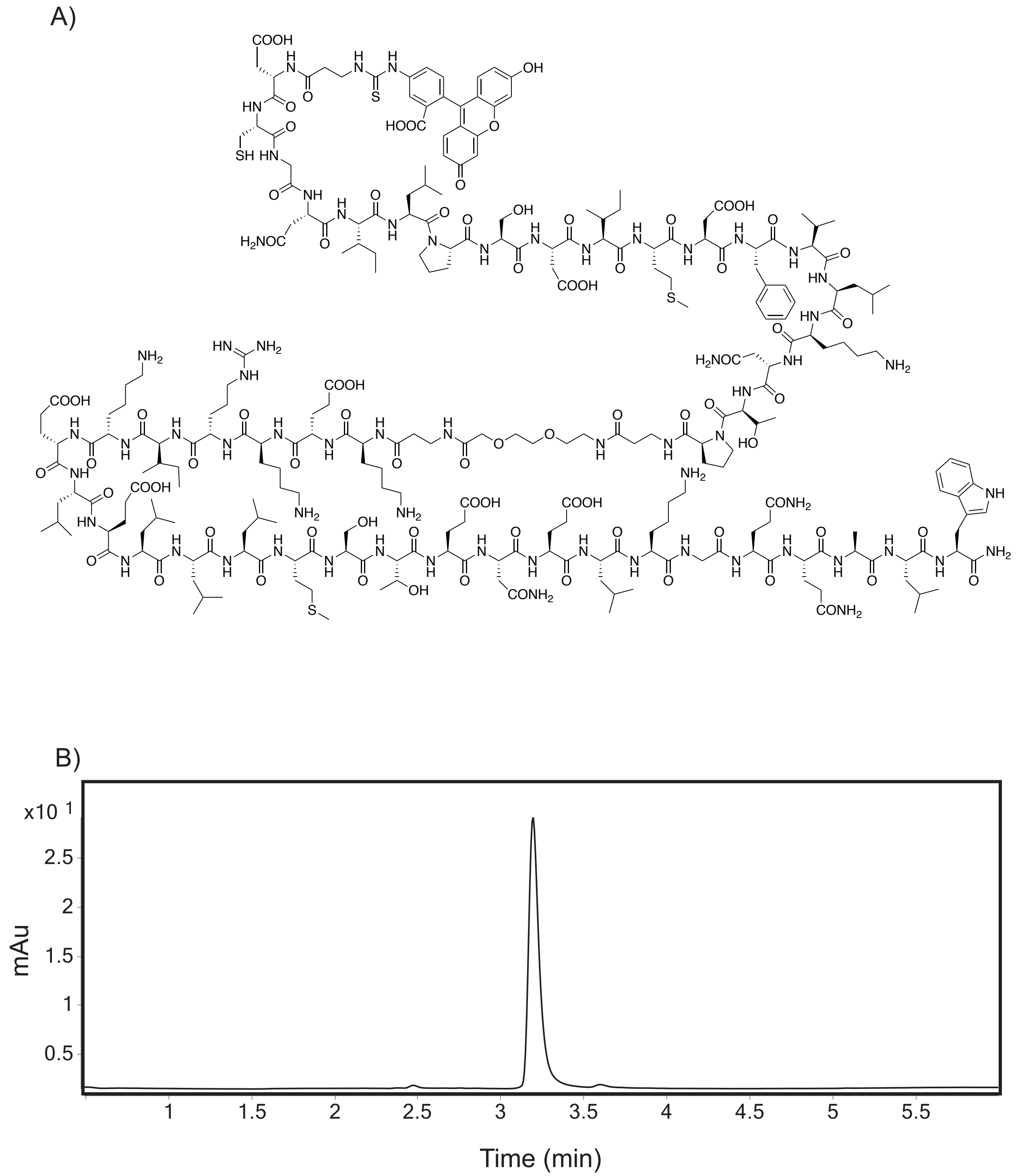


**Figure S10**. A) Structure of **1a**. B) Analytical trace of > 97% purity **1a** at 280 nm.

A)

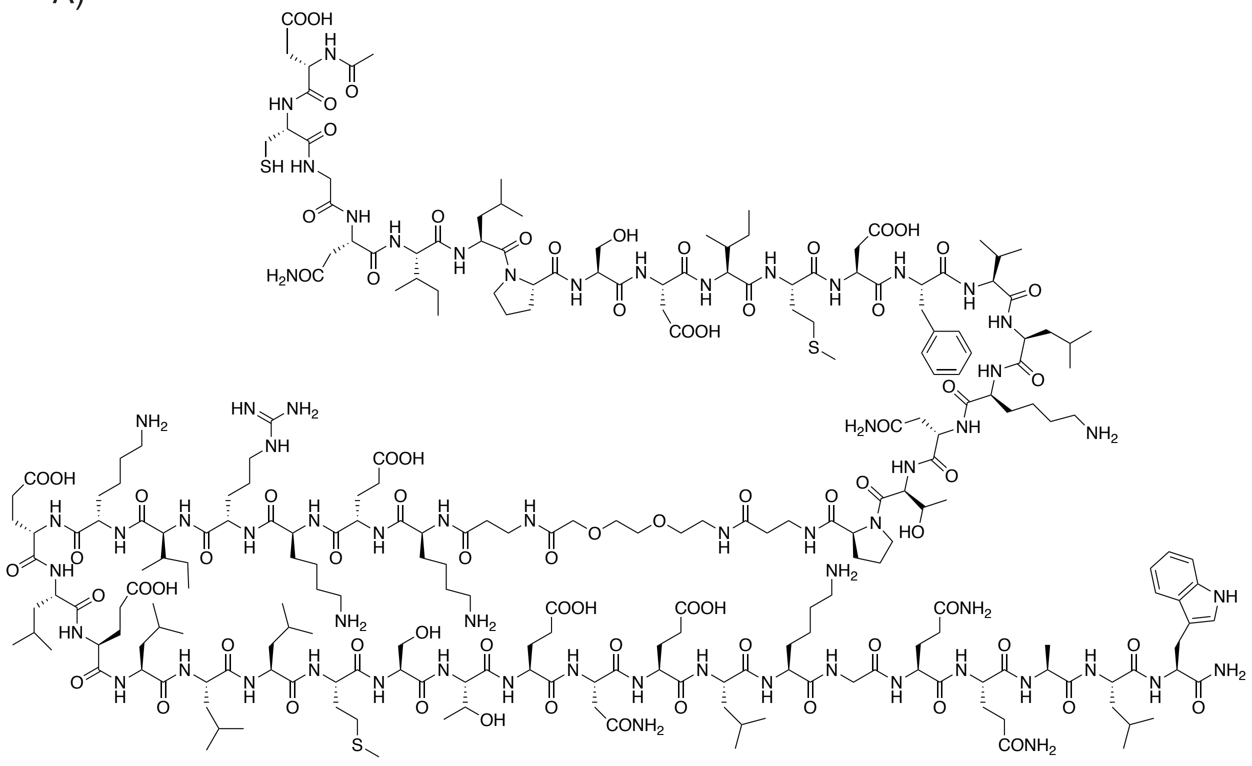

B)

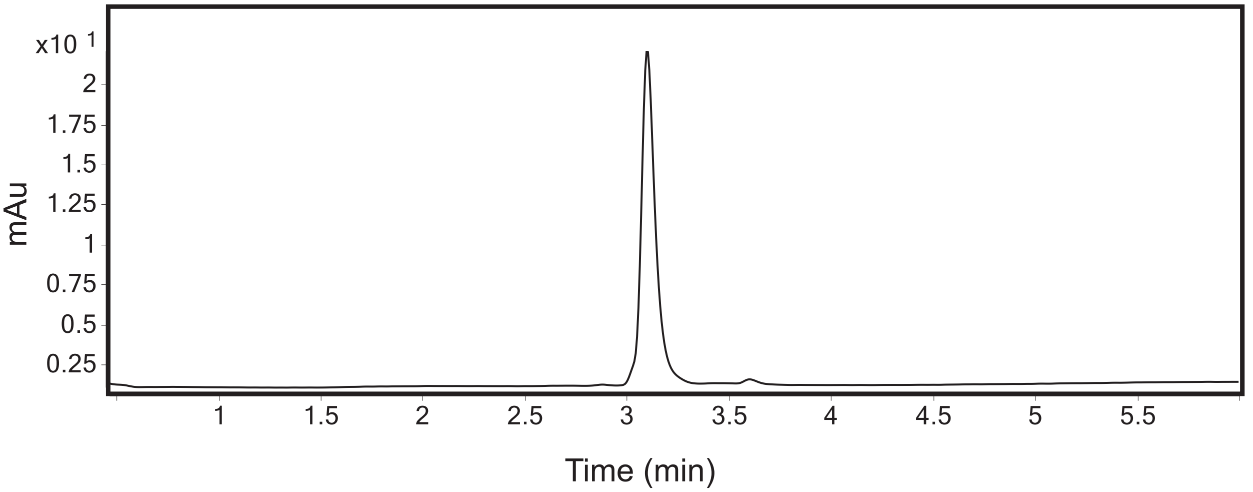


**Figure S11**. A) Structure of **1b**. B) Analytical trace of > 97% purity **1b** at 280 nm.

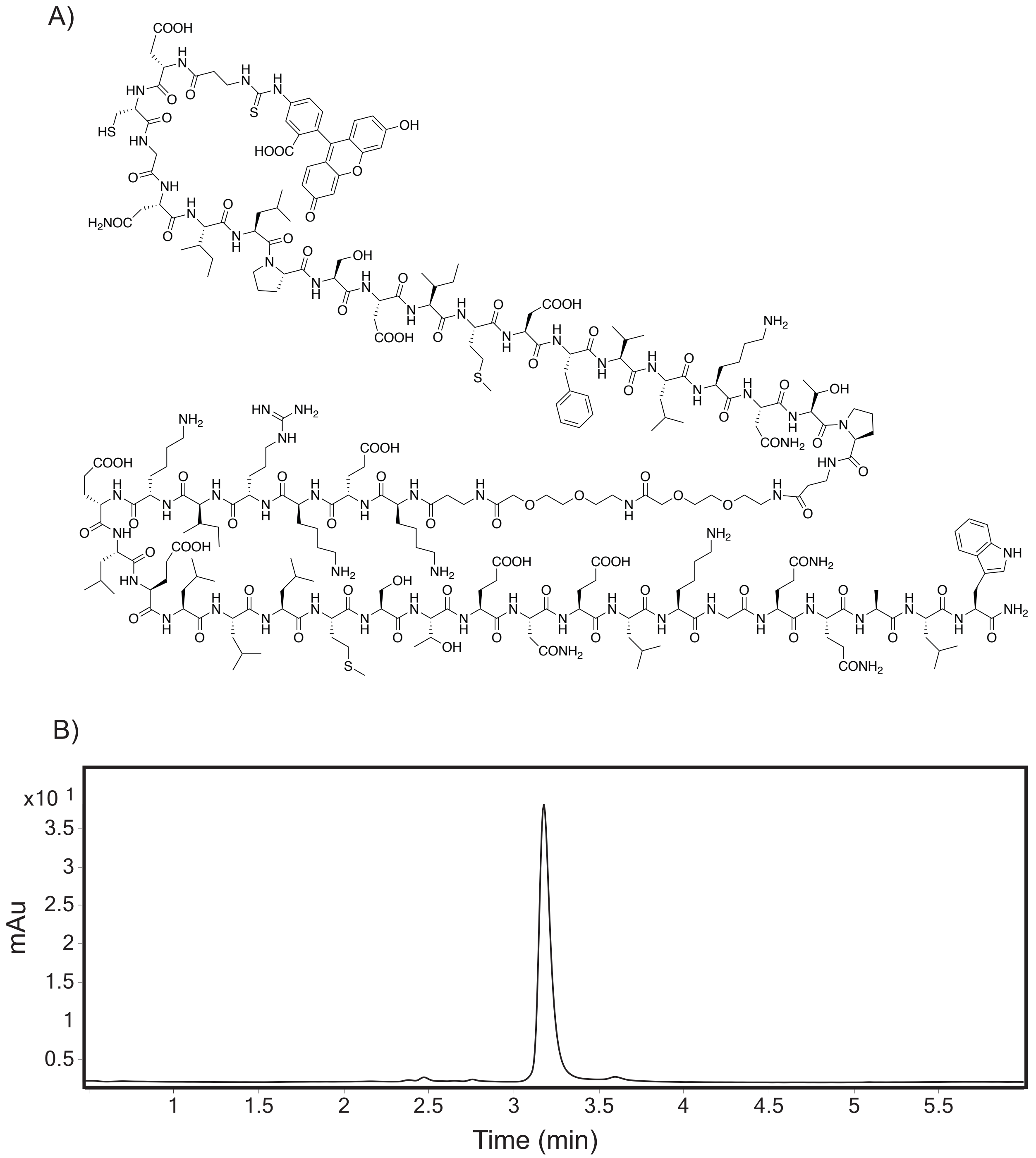


**Figure S12**. A) Structure of **2a**. B) Analytical trace of > 95% purity **2a** at 280 nm.

A)

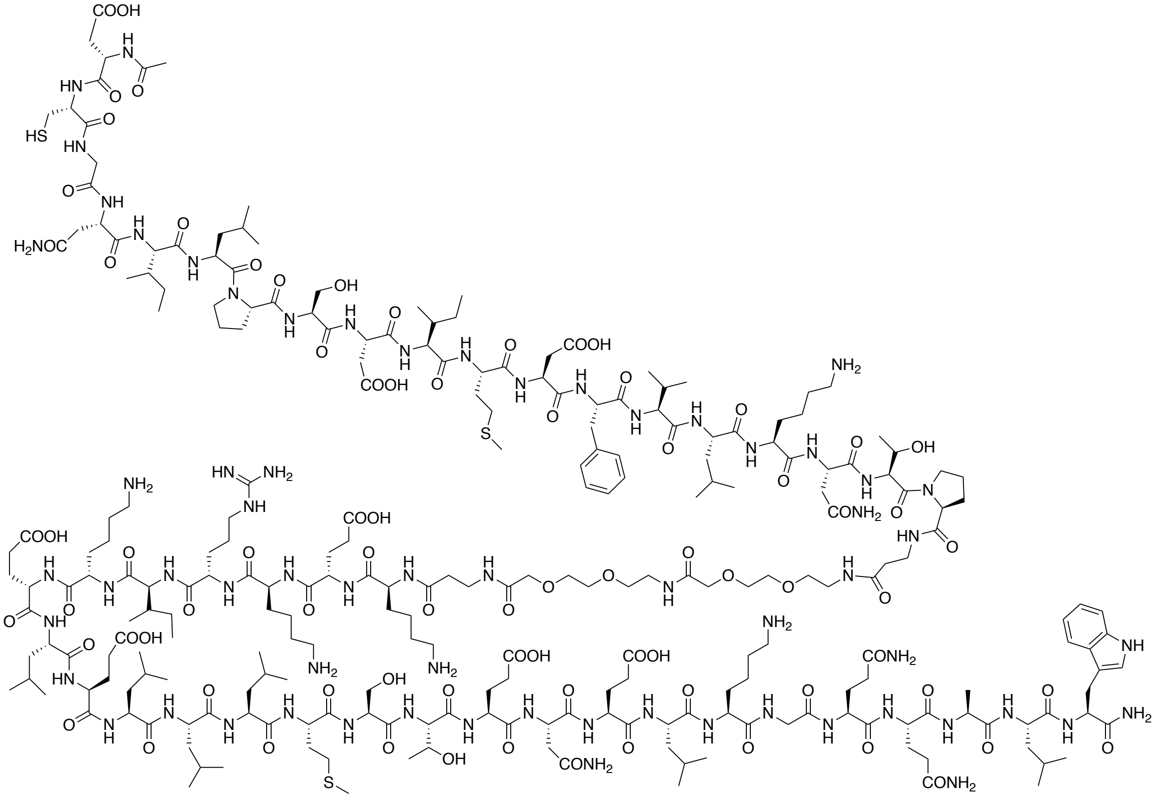

B)

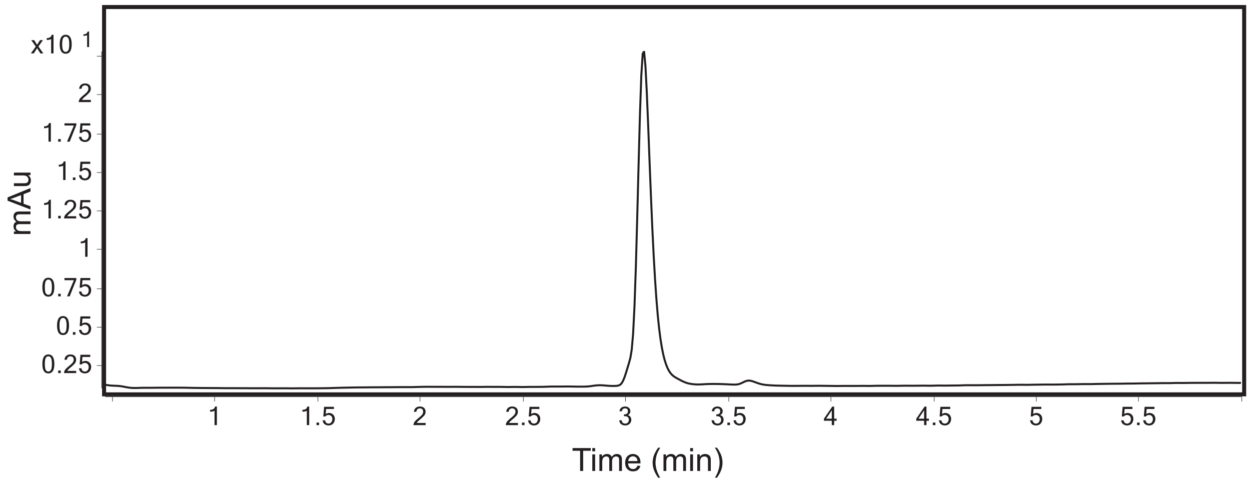


**Figure S13.** A) Structure of **2b**. B) Analytical trace of > 97% purity **2b** at 280 nm.

A)

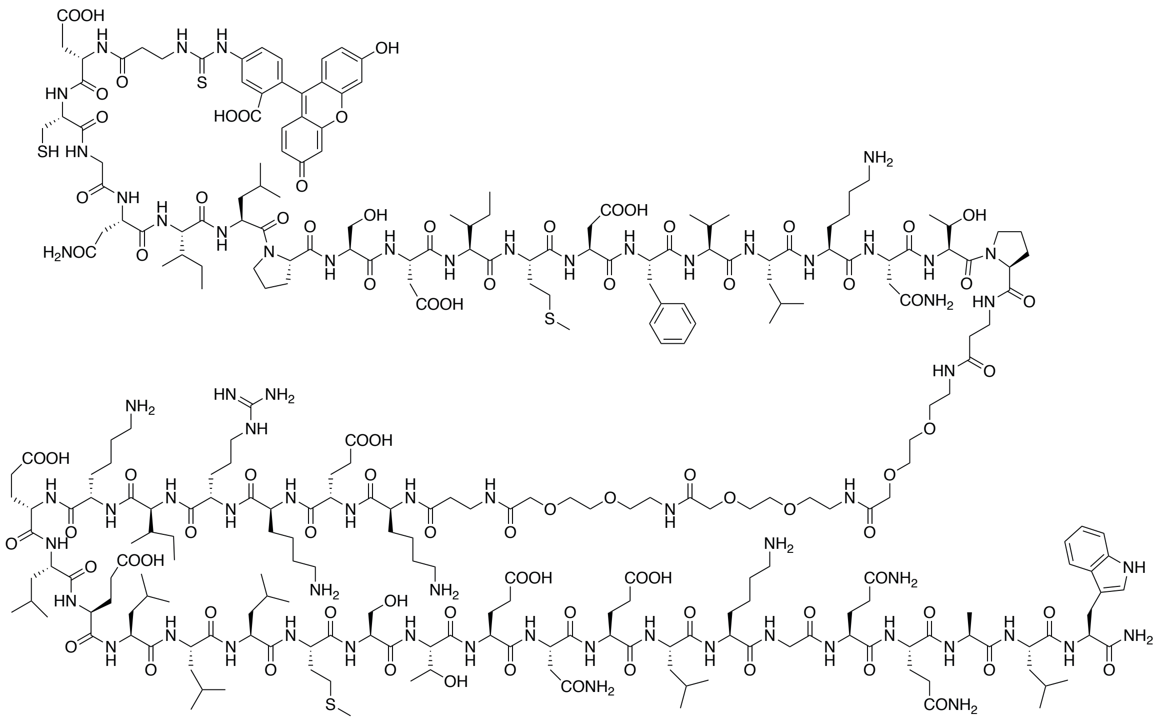

B)

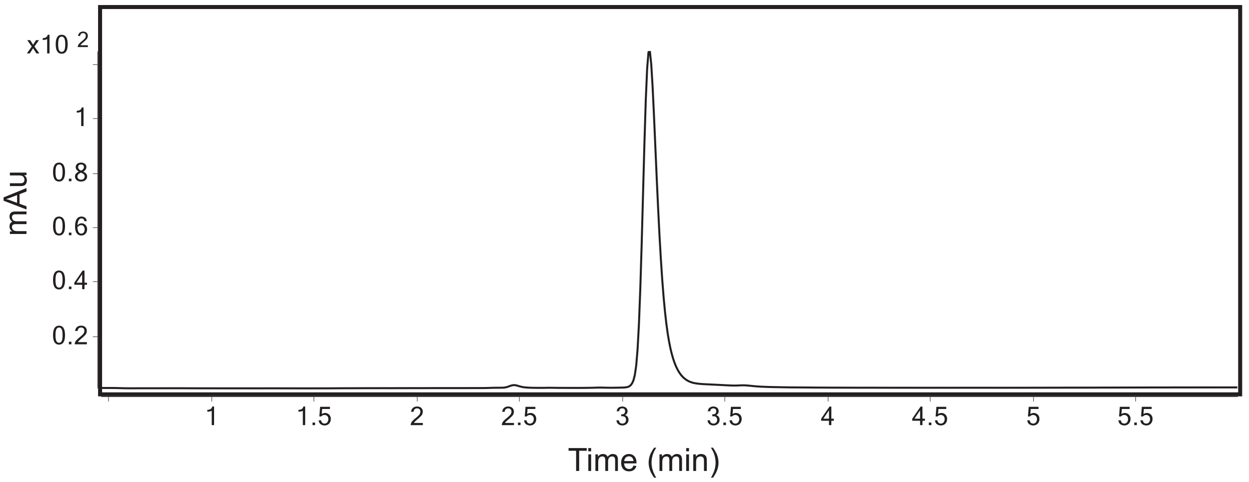


**Figure S14**. A) Structure of **3a**. B) Analytical trace of > 98% purity **3a** at 280 nm.

A)

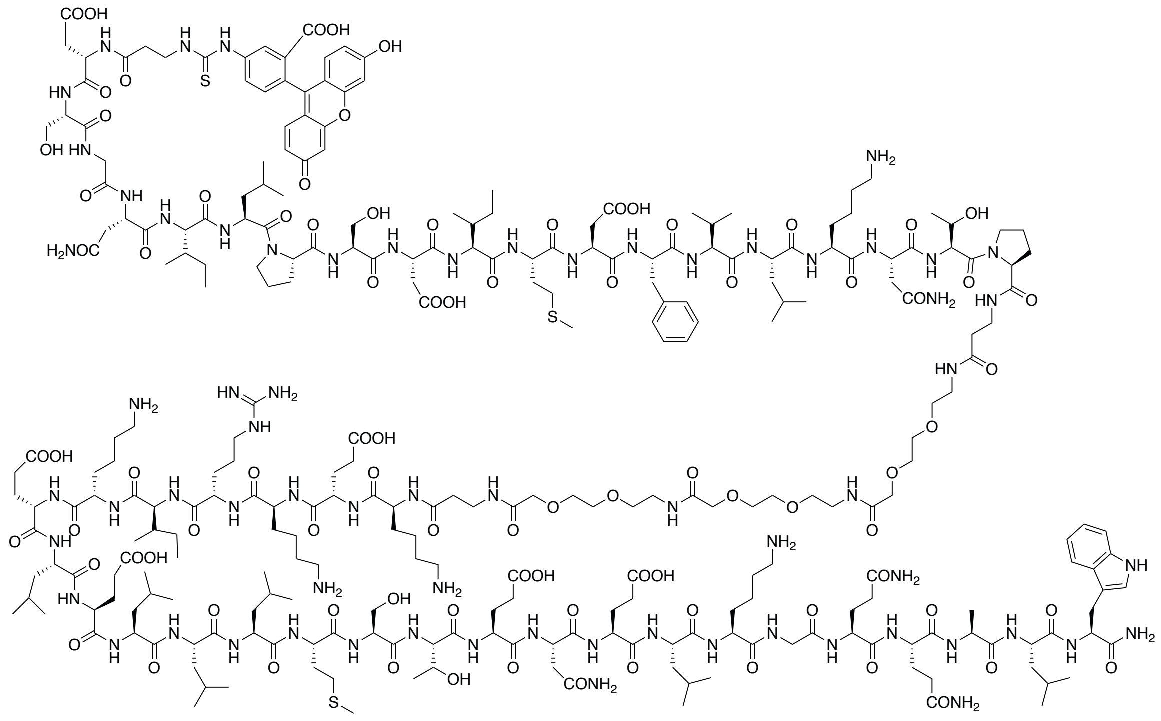

B)

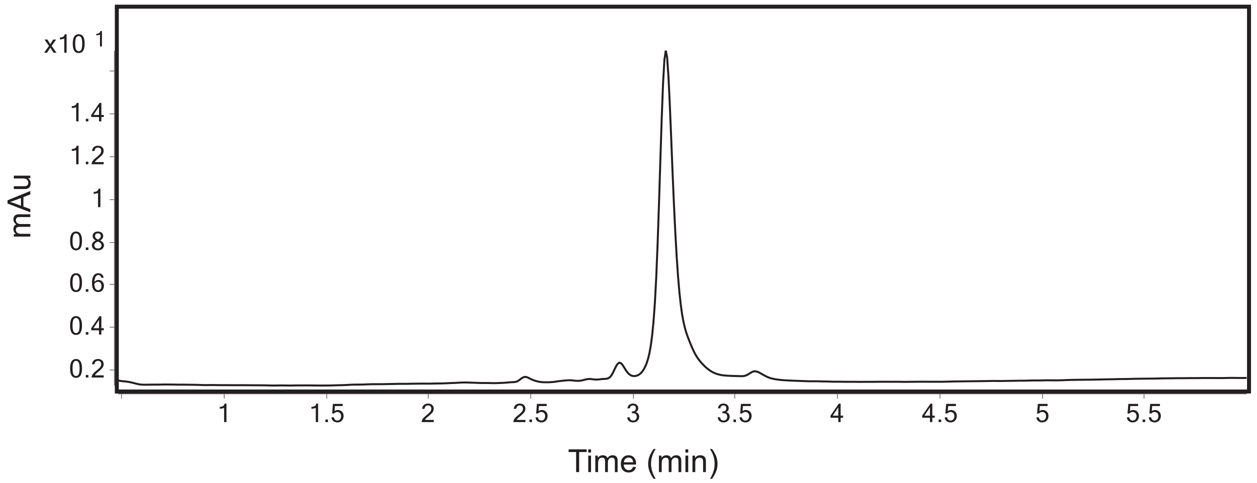


**Figure S15**. A) Structure of **3a** with C841S mutation. B) Analytical trace of > 92% purity C841S **3a** at 280 nm.

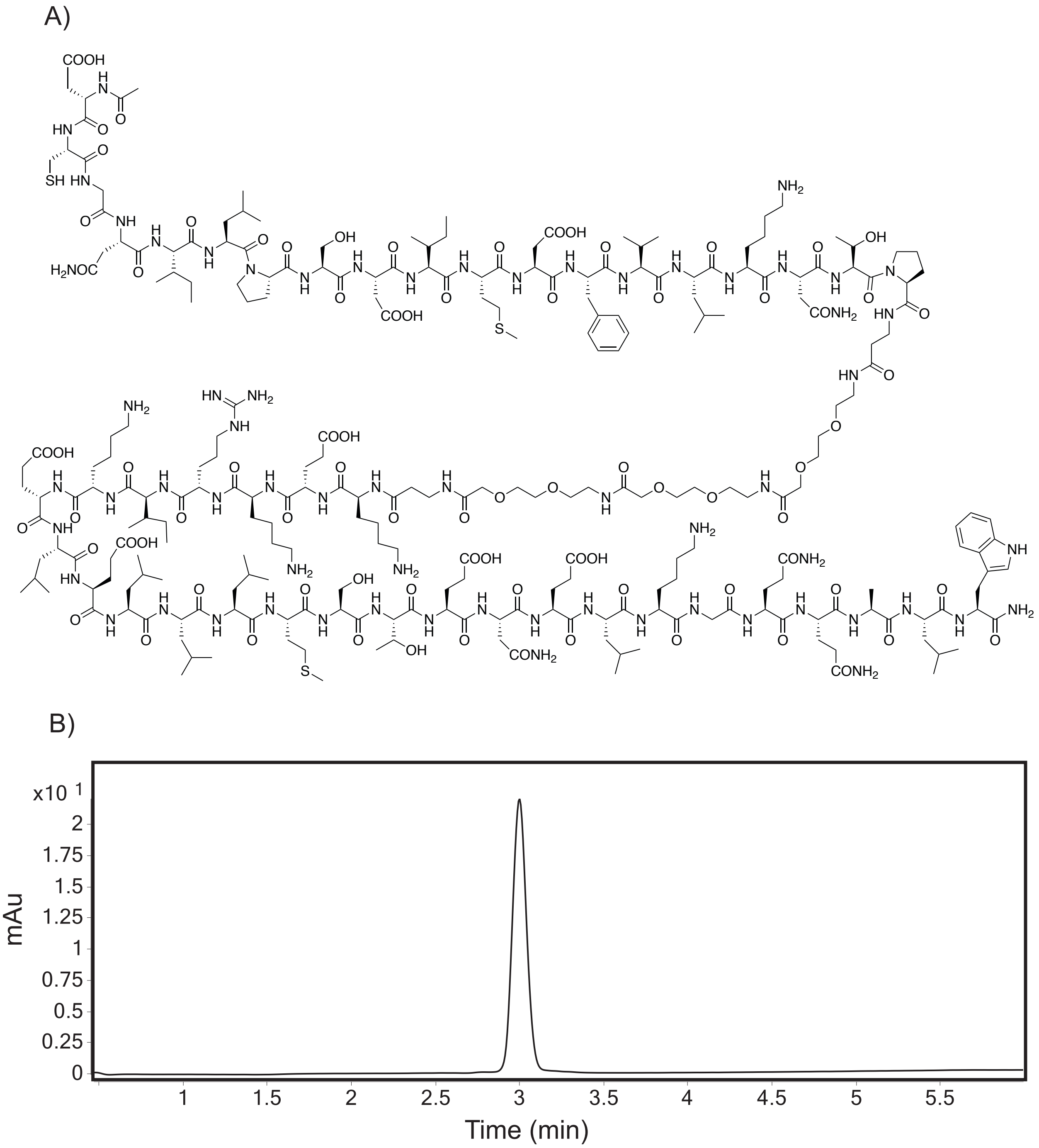


**Figure S16**. A) Structure of **3b**. B) Analytical trace of > 92% purity **3b** at 280 nm.

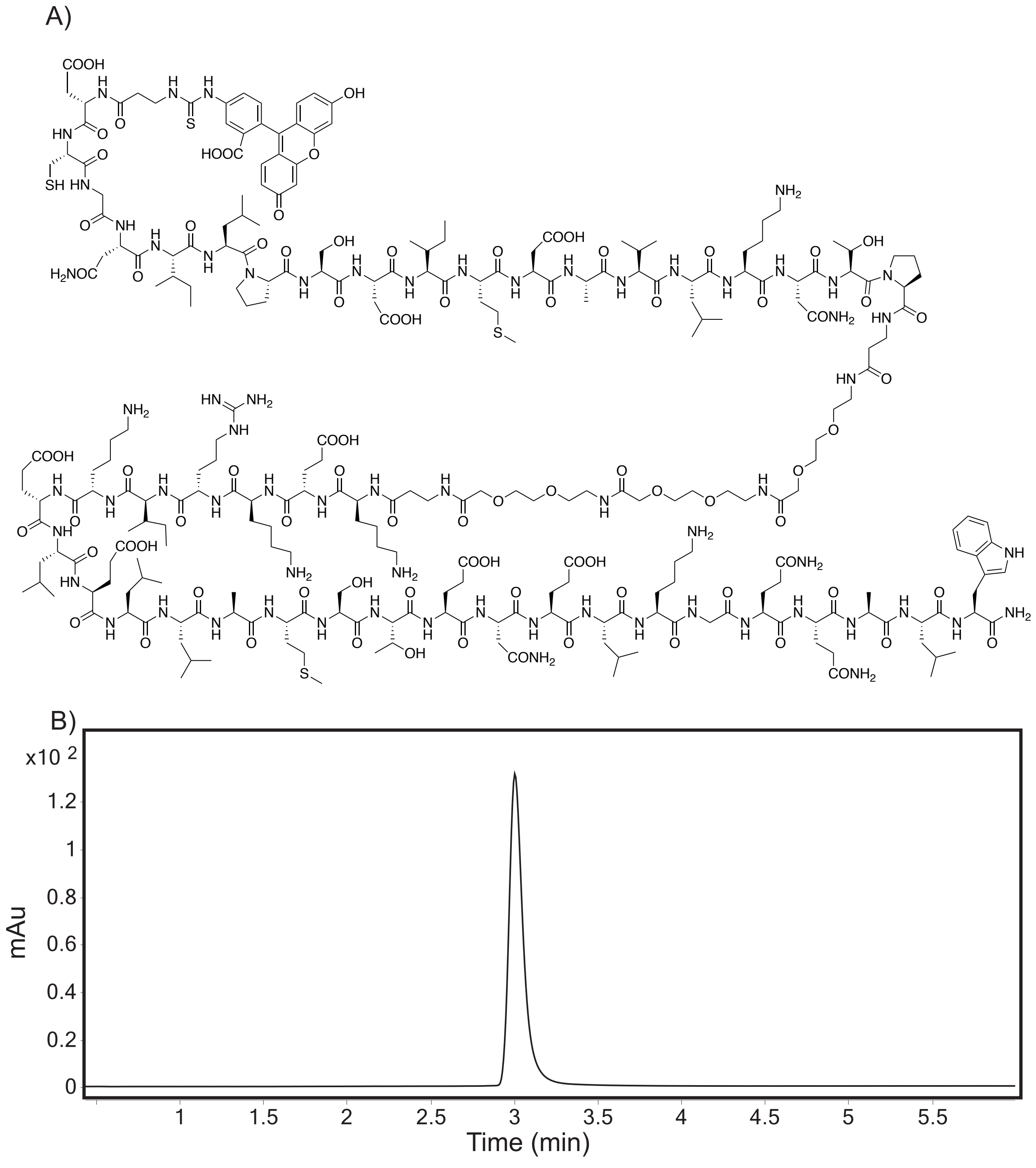


**Figure S17**. A) Structure of **4a**. B) Analytical trace of > 98% purity **4a** at 280 nm.

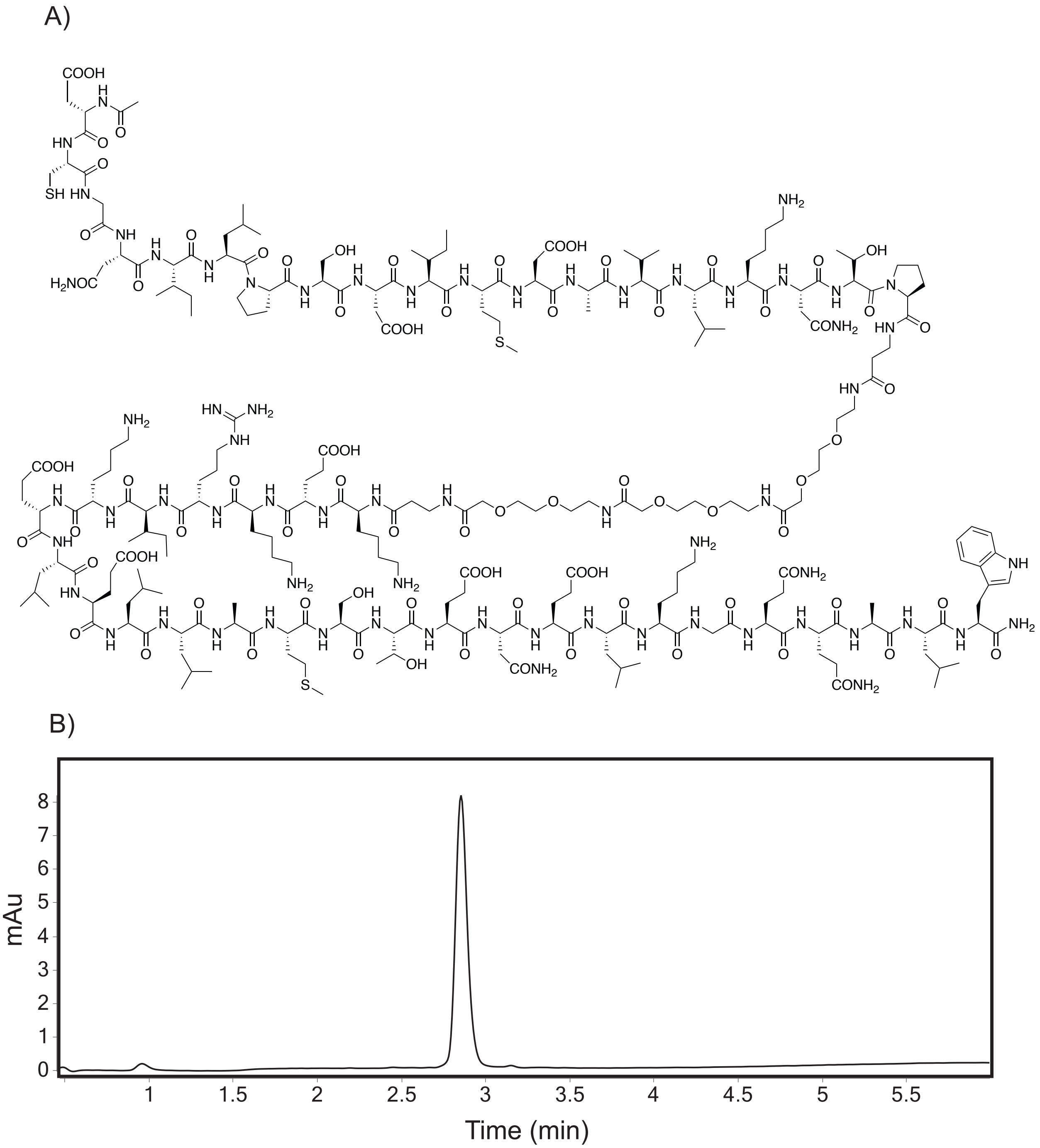


**Figure S18**. A) Structure of **4b**. B) Analytical trace of > 97% purity **4b** at 280 nm.

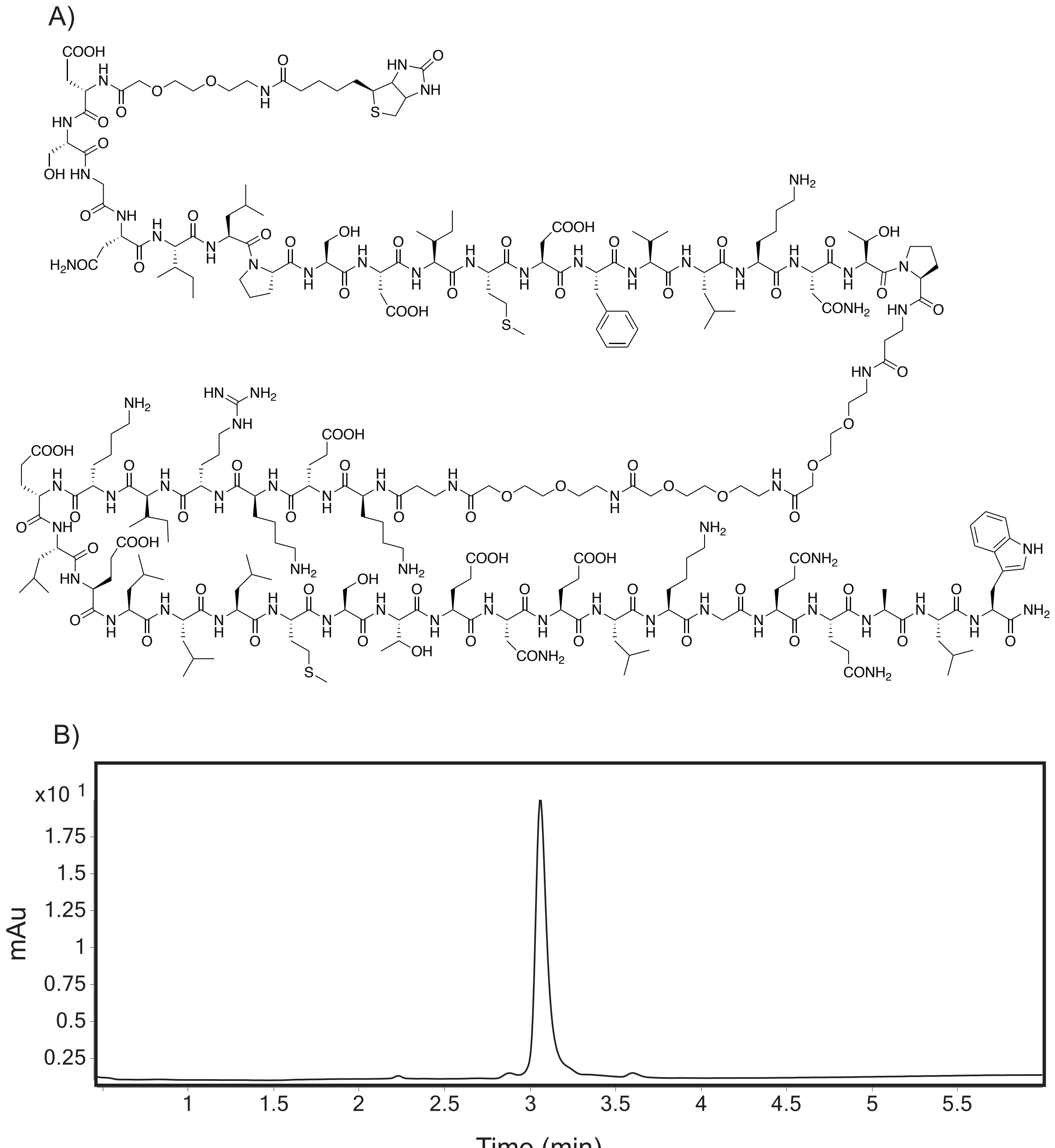

**Figure S19**. A) Structure of **5**. B) Analytical trace of > 96% purity **5** at 280 nm.

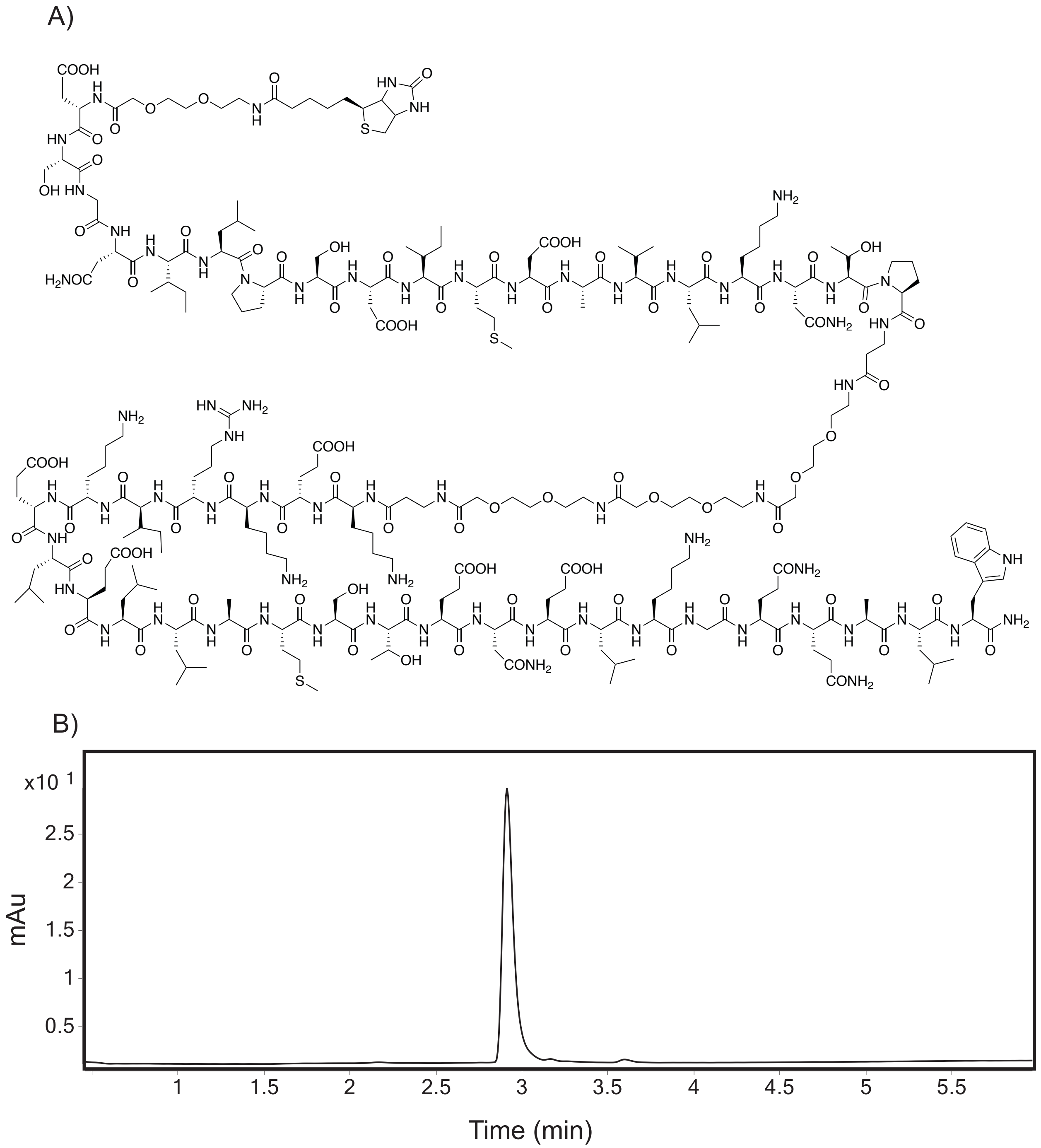


**Figure S20**. A) Structure of **6**. B) Analytical trace of > 96% purity **6** at 280 nm.

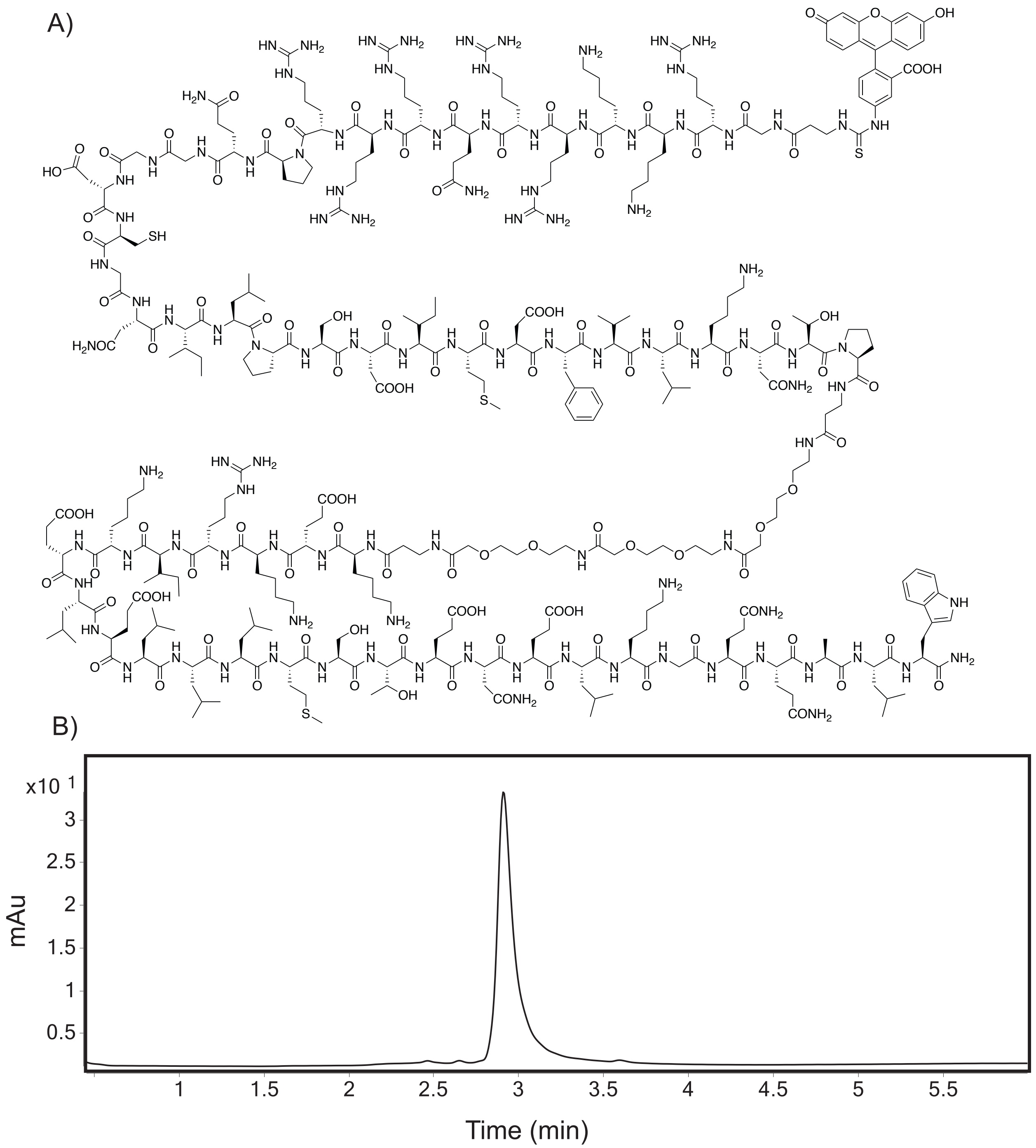


**Figure S21**. A) Structure of **7**. B) Analytical trace of > 93% purity **7** at 280 nm.

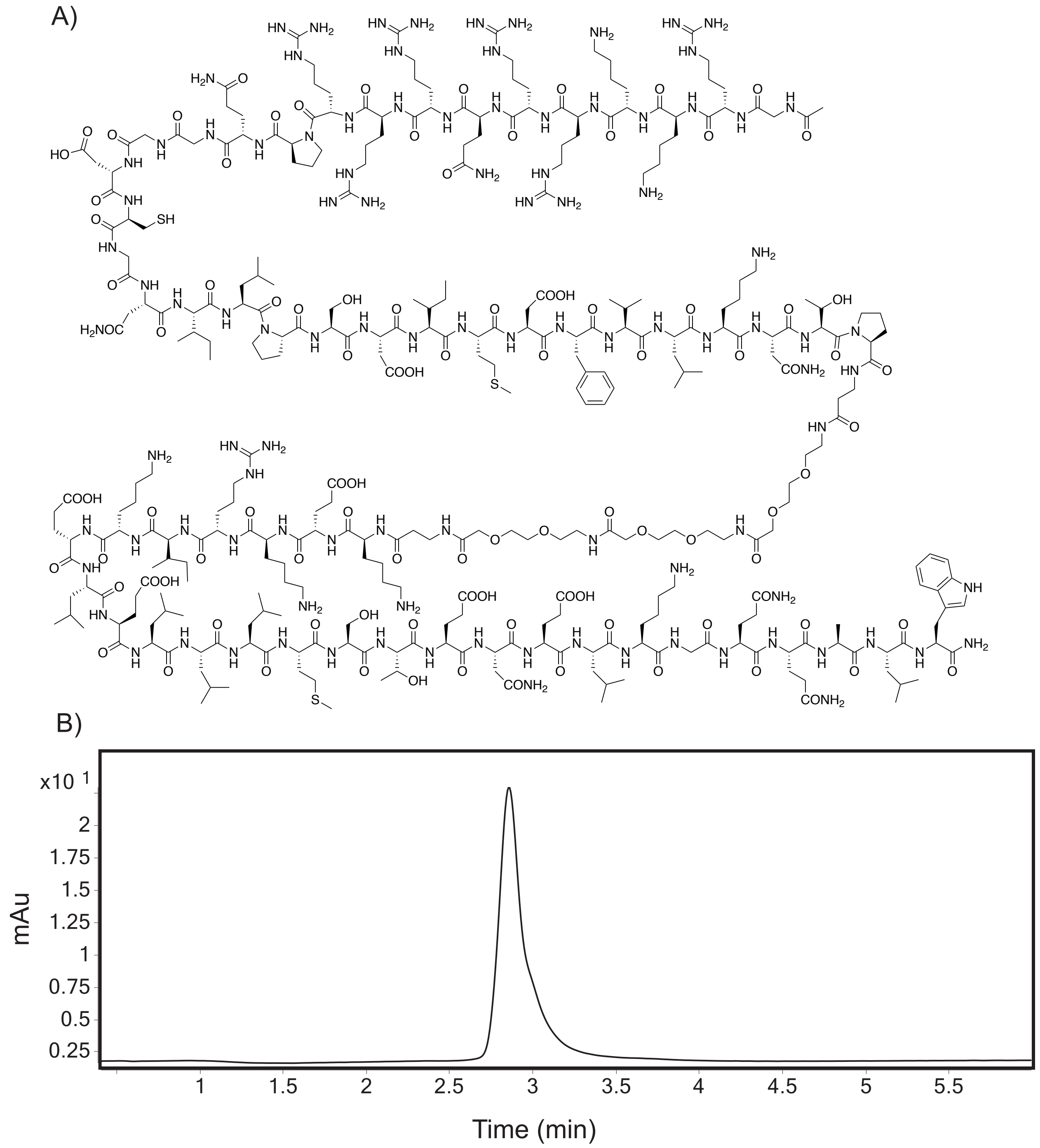


**Figure S22**. A) Structure of **8**. B) Analytical trace of > 97% purity **8** at 280 nm.

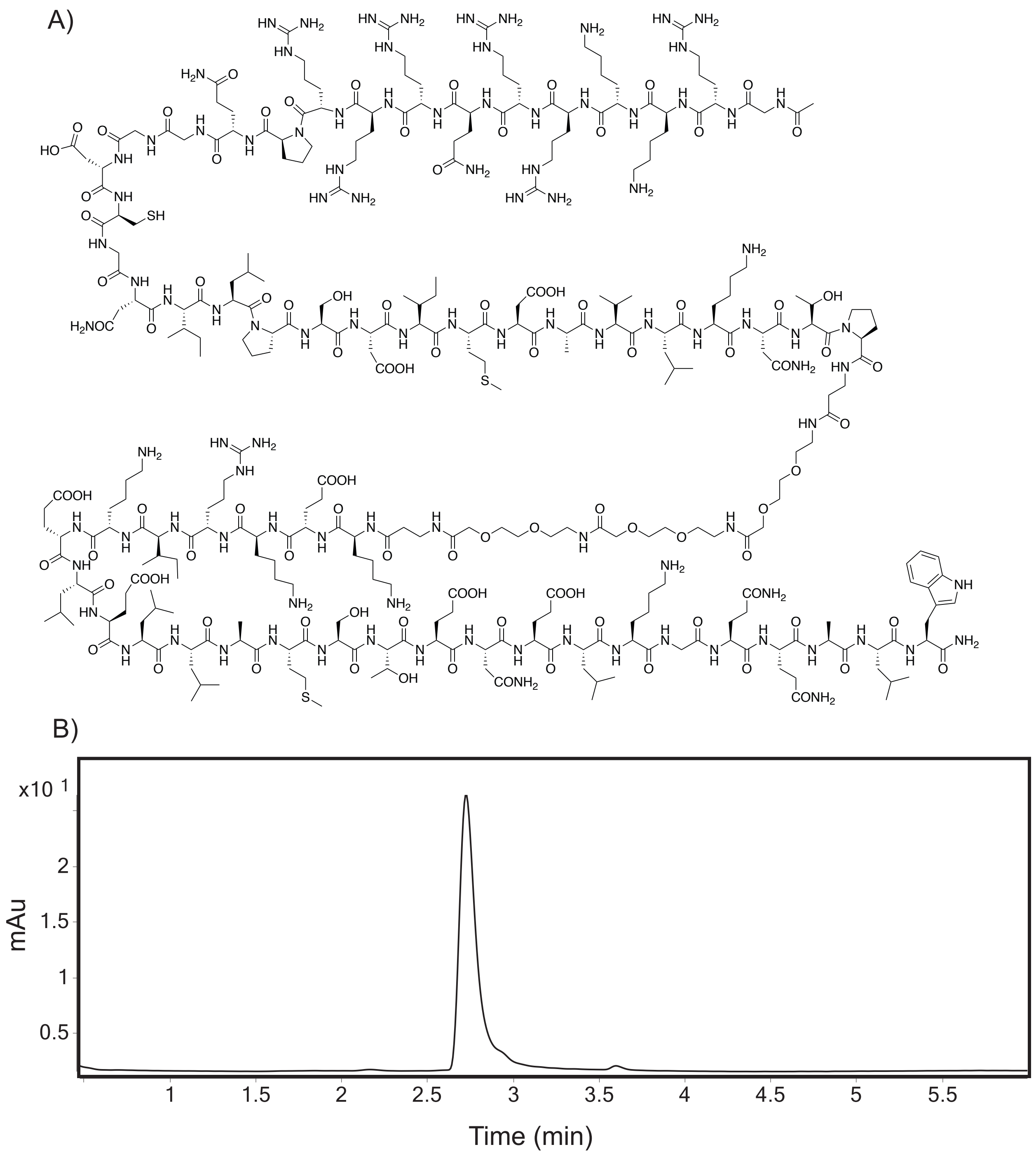


**Figure S23**. A) Structure of **9**. B) Analytical trace of > 97% purity **9** at 280 nm.

A)

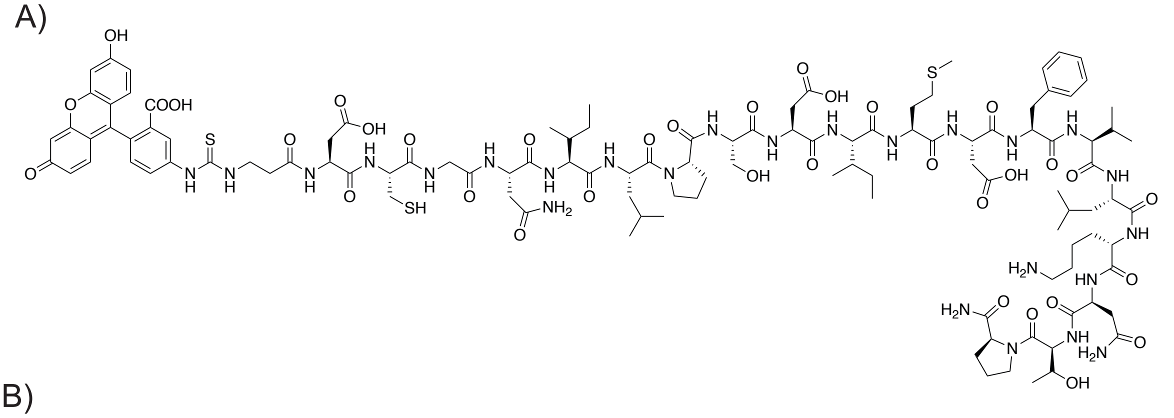

B)

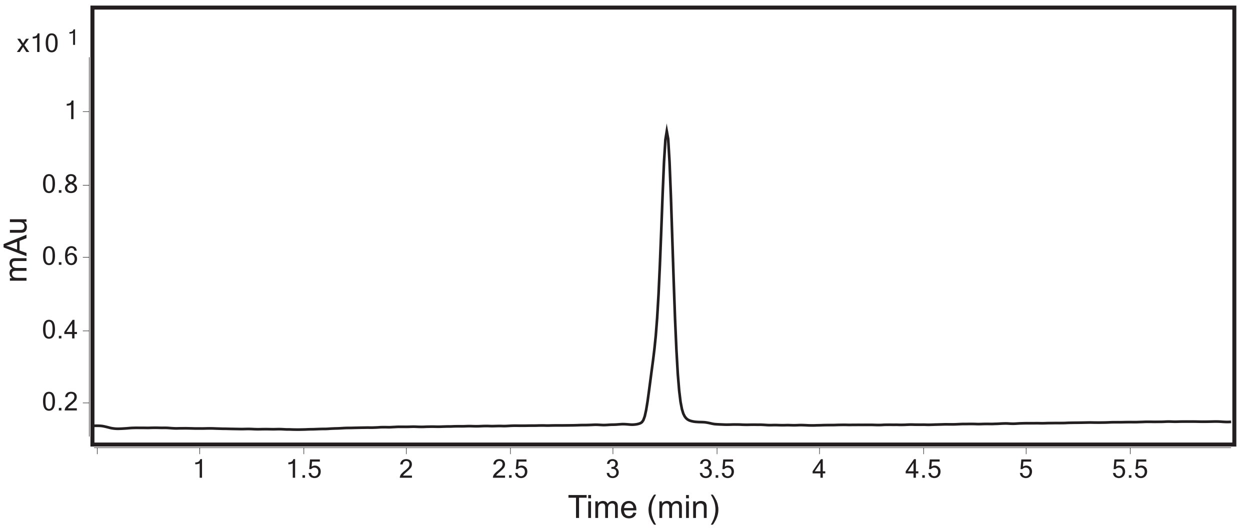


**Figure S24**. A) Structure of FITC-MLL. B) Analytical trace of > 98% purity FITC-MLL at 280 nm.

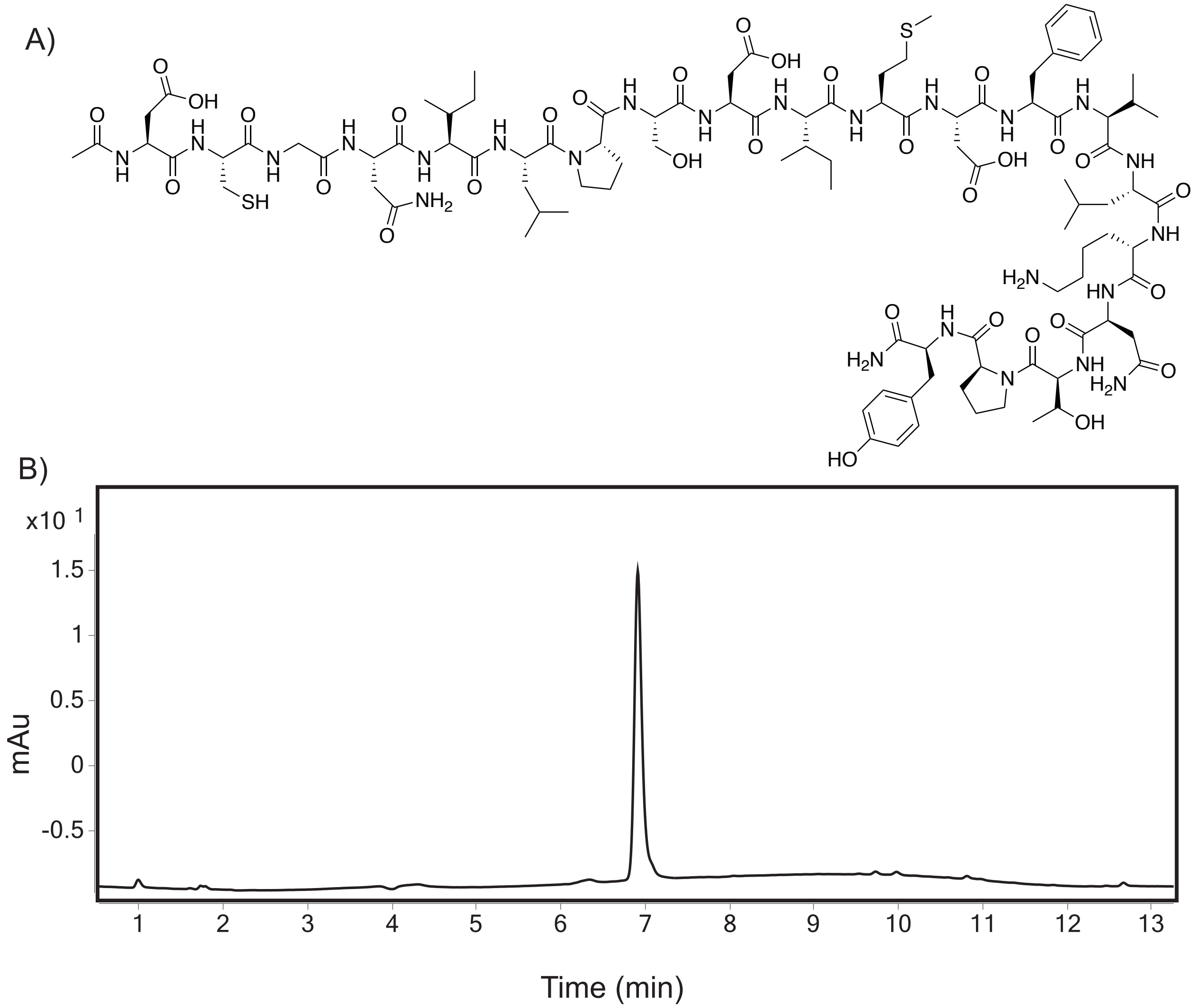

**Figure S25**. A) Structure of MLL. B) Analytical trace of > 96% purity MLL at 280 nm.

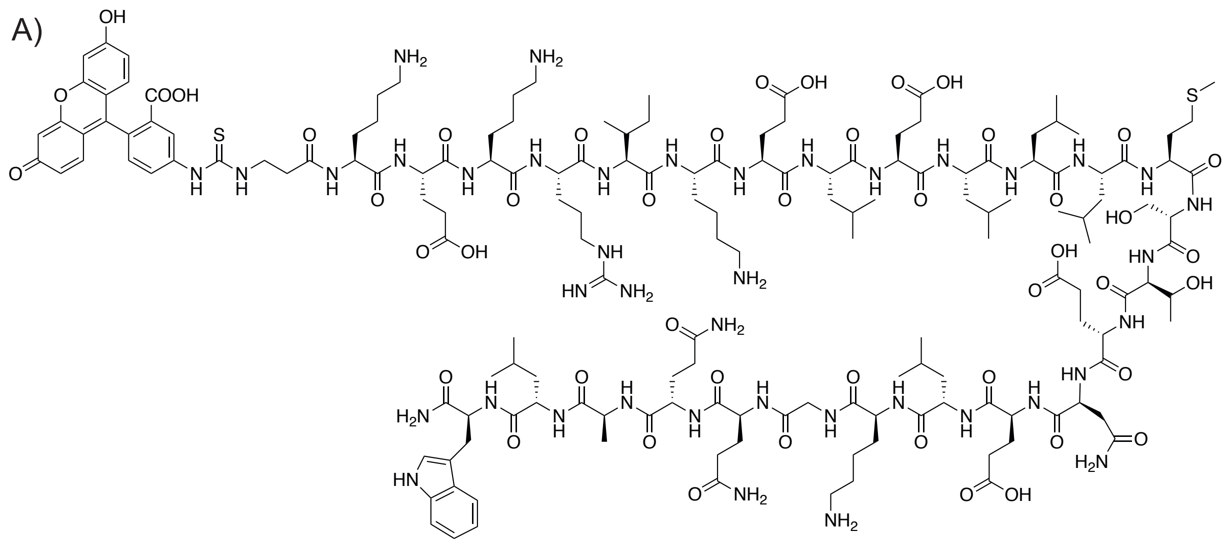

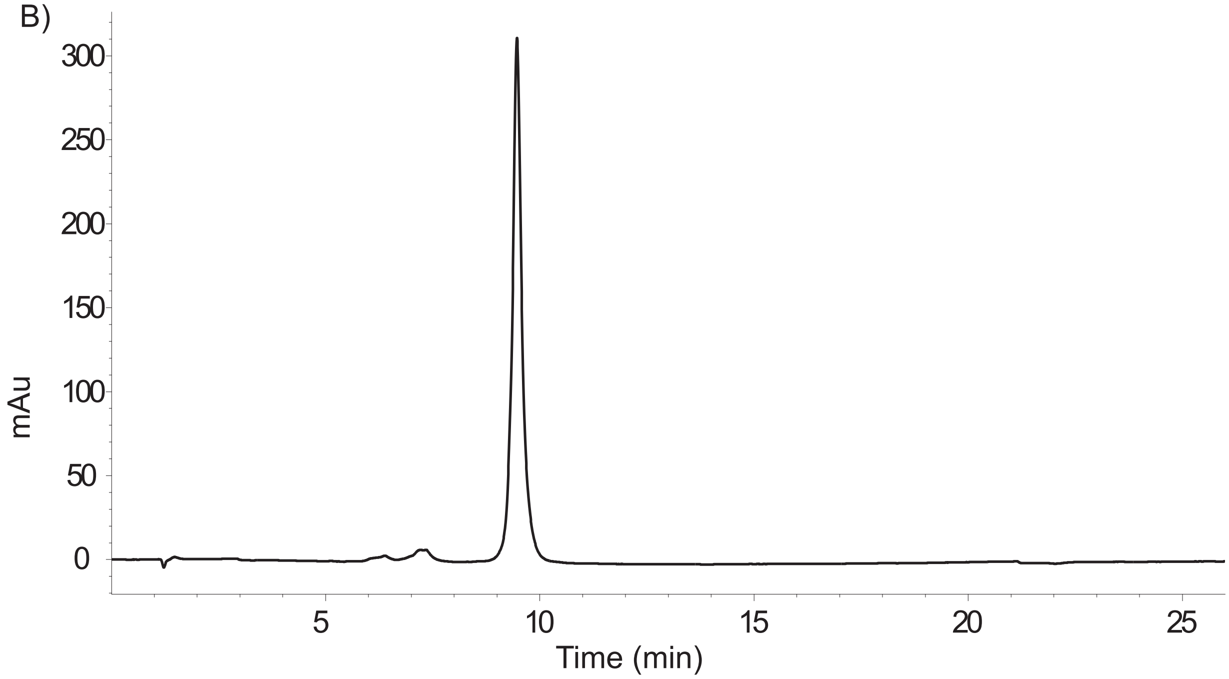


**Figure S26**. A) Structure of FITC-Myb. B) Analytical trace of > 93% purity FITC-Myb at 280 nm.

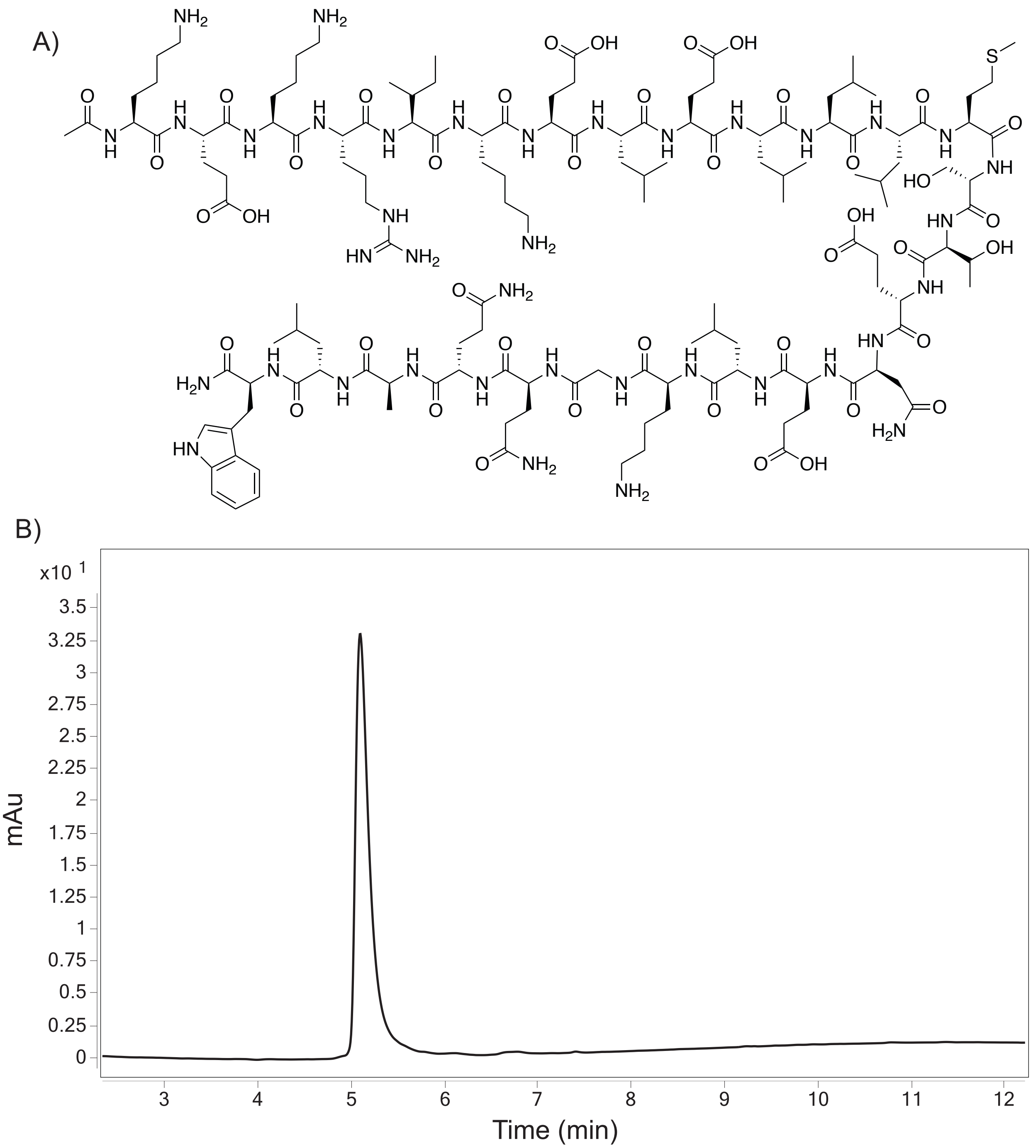


**Figure S27**. A) Structure of Myb. B) Analytical trace of > 98% purity Myb at 280 nm.

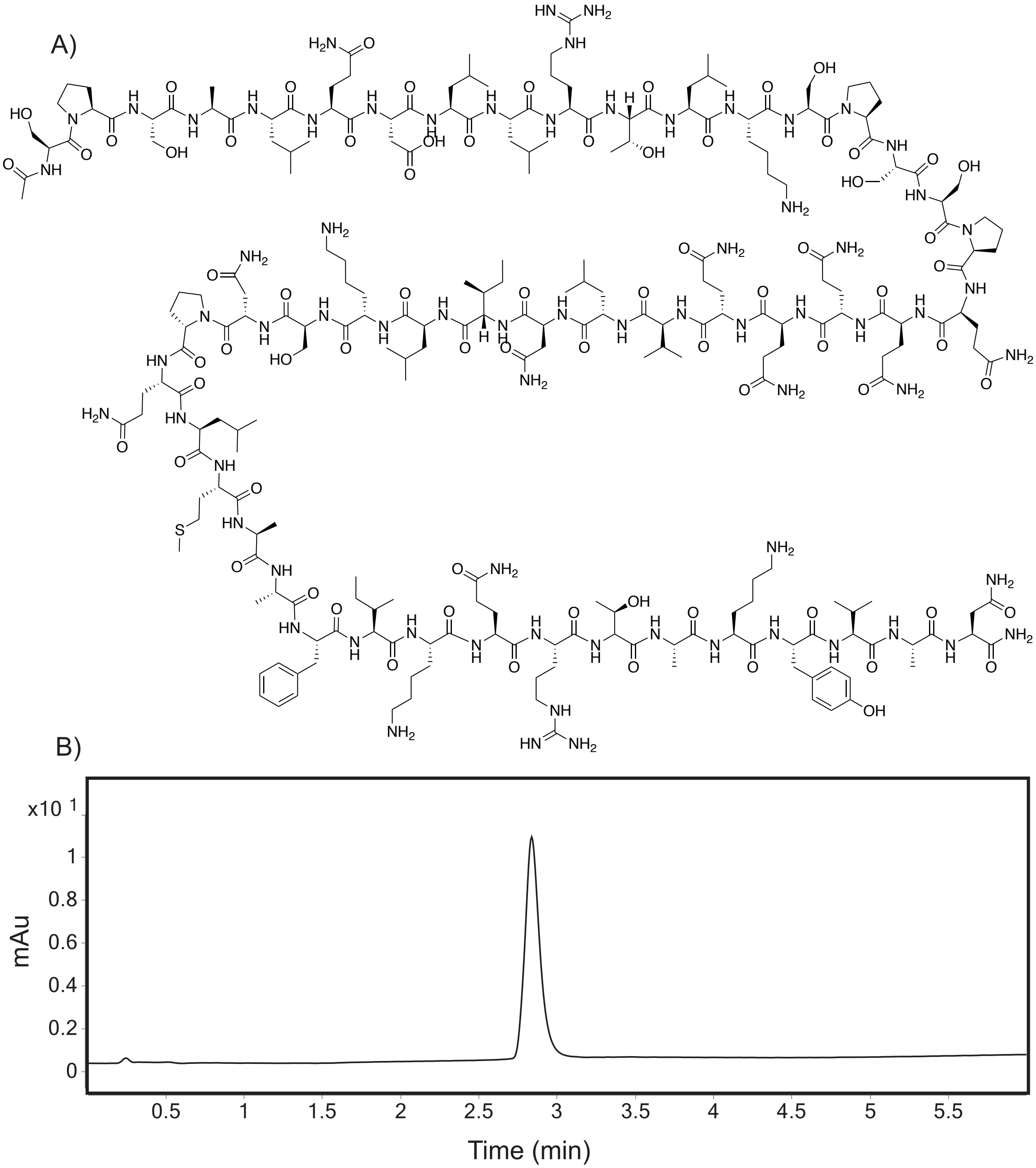


**Figure S28**. A) Structure of CBP IBiD (2063-2111). B) Analytical trace of > 98% purity CBP IBiD (2063-2111) at 280 nm.

**References**.


(1) Buhrlage, S. J.; Bates, C. A.; Rowe, S. P.; Minter, A. R.; Brennan, B. B.; Majmudar, C. Y.; Wemmer, D. E.; Al-Hashimi, H.; Mapp, A. K. Amphipathic Small Molecules Mimic the Binding Mode and Function of Endogenous Transcription Factors. *ACS Chem. Biol.* **2009**, *4* (5), 335–344. https://doi.org/10.1021/cb900028j.

(2) Thakur, J. K.; Arthanari, H.; Yang, F.; Pan, S.-J.; Fan, X.; Breger, J.; Frueh, D. P.; Gulshan, K.; Li, D. K.; Mylonakis, E.; Struhl, K.; Moye-Rowley, W. S.; Cormack, B. P.; Wagner, G.; Näär, A. M. A Nuclear Receptor-like Pathway Regulating Multidrug Resistance in Fungi. *Nature* **2008**, *452* (7187), 604–609. https://doi.org/10.1038/nature06836.

(3) Pomerantz, W. C.; Wang, N.; Lipinski, A. K.; Wang, R.; Cierpicki, T.; Mapp, A. K. Profiling the Dynamic Interfaces of Fluorinated Transcription Complexes for Ligand Discovery and Characterization. *ACS Chem. Biol.* **2012**, *7* (8), 1345–1350. https://doi.org/10.1021/cb3002733.

(4) Yang, F.; Vought, B. W.; Satterlee, J. S.; Walker, A. K.; Jim Sun, Z.-Y.; Watts, J. L.; DeBeaumont, R.; Mako Saito, R.; Hyberts, S. G.; Yang, S.; Macol, C.; Iyer, L.; Tjian, R.; van den Heuvel, S.; Hart, A. C.; Wagner, G.; Näär, A. M. An ARC/Mediator Subunit Required for SREBP Control of Cholesterol and Lipid Homeostasis. *Nature* **2006**, *442* (7103), 700–704. https://doi.org/10.1038/nature04942.

(5) Nagulapalli, M.; Maji, S.; Dwivedi, N.; Dahiya, P.; Thakur, J. K. Evolution of Disorder in Mediator Complex and Its Functional Relevance. *Nucleic Acids Res.* **2016**, *44* (4), 1591–1612. https://doi.org/10.1093/nar/gkv1135.

(6) Nishikawa, J. L.; Boeszoermenyi, A.; Vale-Silva, L. A.; Torelli, R.; Posteraro, B.; Sohn, Y.-J.; Ji, F.; Gelev, V.; Sanglard, D.; Sanguinetti, M.; Sadreyev, R. I.; Mukherjee, G.; Bhyravabhotla, J.; Buhrlage, S. J.; Gray, N. S.; Wagner, G.; Näär, A. M.; Arthanari, H. Inhibiting Fungal Multidrug Resistance by Disrupting an Activator–Mediator Interaction. *Nature* **2016**, *530* (7591), 485–489. https://doi.org/10.1038/nature16963.

(7) Vojnic, E.; Mourão, A.; Seizl, M.; Simon, B.; Wenzeck, L.; Larivière, L.; Baumli, S.; Baumgart, K.; Meisterernst, M.; Sattler, M.; Cramer, P. Structure and VP16 Binding of the Mediator Med25 Activator Interaction Domain. *Nat. Struct. Mol. Biol.* **2011**, *18* (4), 404–409. https://doi.org/10.1038/nsmb.1997.

(8) Bontems, F.; Verger, A.; Dewitte, F.; Lens, Z.; Baert, J.-L.; Ferreira, E.; Launoit, Y. de; Sizun, C.; Guittet, E.; Villeret, V.; Monté, D. NMR Structure of the Human Mediator MED25 ACID Domain. *J. Struct. Biol.* **2011**, *174* (1), 245–251. https://doi.org/10.1016/j.jsb.2010.10.011.

(9) Majmudar, C. Y.; Højfeldt, J. W.; Arevang, C. J.; Pomerantz, W. C.; Gagnon, J. K.; Schultz, P. J.; Cesa, L. C.; Doss, C. H.; Rowe, S. P.; Vásquez, V.; Tamayo-Castillo, G.; Cierpicki, T.; Brooks, C. L.; Sherman, D. H.; Mapp, A. K. Sekikaic Acid and Lobaric Acid Target a Dynamic Interface of the Coactivator CBP/P300. *Angew. Chem. Int. Ed.* **2012**, *51* (45), 11258–11262. https://doi.org/10.1002/anie.201206815.

(10) Roehrl, M. H. A.; Wang, J. Y.; Wagner, G. A General Framework for Development and Data Analysis of Competitive High-Throughput Screens for Small-Molecule Inhibitors of Protein−Protein Interactions by Fluorescence Polarization †. *Biochemistry* **2004**, *43* (51), 16056–16066. https://doi.org/10.1021/bi048233g.

(11) Malatesta, F. The Study of Bimolecular Reactions under Non-Pseudo-First Order Conditions. *Biophys. Chem.* **2005**, *116* (3), 251–256. https://doi.org/10.1016/j.bpc.2005.04.006.

(12) Rogers, J. M.; Steward, A.; Clarke, J. Folding and Binding of an Intrinsically Disordered Protein: Fast, but Not 'Diffusion-Limited.' *J. Am. Chem. Soc.* **2013**, *135* (4), 1415–1422. https://doi.org/10.1021/ja309527h.
(13) De Guzman, R. N.; Goto, N. K.; Dyson, H. J.; Wright, P. E. Structural Basis for Cooperative Transcription Factor Binding to the CBP Coactivator. *J. Mol. Biol.* **2006**, *355* (5), 1005–1013. https://doi.org/10.1016/j.jmb.2005.09.059.
